# Structures of MyD88 and its oncogenic mutant reveal DNA binding with implications for cytosolic DNA sensing

**DOI:** 10.64898/2026.10.21.760499

**Authors:** Pietro Fontana, Ahmet B. Tufan, Patrick Günther, Elena Goldberg, Sebastian Dörfler, Setu M. Vora, Leo Schirokauer, Joao A. Paulo, Hyung Suk Oh, Steven P. Gygi, David M. Knipe, Hao Wu

**Author notes:** These authors contributed equally (Pietro Fontana, Ahmet B. Tufan, Patrick Günther). Department of Immunology, St. Jude Children’s Research Hospital, Memphis, TN.

## Abstract

MyD88 is a universal adaptor that connects to Toll-like receptors and interleukin-1 receptors through its Toll/IL-1R homology (TIR) domain and activates inflammatory signaling via its death domain (DD). However, how MyD88^TIR^ serves as the platform for these functions and if it has additional functions remain unclear. Here, we report cryogenic electron microscopy (cryo-EM) structures of MyD88^TIR^ and its cancer-specific mutant L252P (previously L265P) mainly in ring and spiral forms, respectively. Both assemblies are formed by antiparallel double-filaments, using canonical TIR interfaces within each strand but unique surfaces between the strands, and position MyD88^DD^ at the same sides of the assemblies for self-oligomerization and pathway activation. Surprisingly, these WT or L252P filaments are bound to dsDNA using the concave sides of the rings and the spirals. In vitro assays and cellular studies establish that MyD88 directly binds dsDNA and promotes inflammatory cytokine expression, particularly pro-IL-1β, following dsDNA transfection or viral infection. Notably, these responses are unaffected by deficiency of UNC93B1, the essential chaperone required for trafficking endosomal nucleic acid-sensing TLRs. Structure-guide MyD88 mutant that disrupts dsDNA binding specifically compromises dsDNA-induced transcriptional program while maintaining TLR signaling. RNA-seq further reveals that MyD88 plays a crucial role in inducing gene expression upon dsDNA sensing and that about half of cGAS-dependent genes are also MyD88-dependent. Together, our structural, biochemical and genetic data support the findings that MyD88 directly engages with cytosolic dsDNA and cooperates with cGAS to promote the full innate immune transcriptional response.

## Main

Toll-like receptors (TLRs) and interleukin-1 receptors (IL-1Rs) are key players in the innate immune system.^1,2^ TLRs recognize pathogen-associated molecular patterns (PAMPs), while IL-1Rs detect cytokines from the IL-1 family. Despite differences in their extracellular domains, both receptors share a conserved cytosolic Toll/IL-1R homology (TIR) domain. Upon activation by PAMPs or cytokines, receptors dimerize or oligomerize, which brings the TIR domains into close proximity, creating a platform for recruiting adaptor proteins like TIRAP (also known as Mal), TRIF, TRAM, and myeloid differentiation primary response 88 (MyD88).^1,2^ The adaptor protein MyD88 is particularly well known for its broad involvement in TLR signaling.^1–7^ It is recruited in response to a wide array of activating signals for all TLRs except TLR3.^8,9^ MyD88 also plays a crucial role in mediating the signaling pathways activated by cytokines IL-1, IL-18, and IL-33, which are recognized by the IL-1R family.^10^

MyD88 consists of two distinct domains separated by a linker: an N-terminal death domain (DD) and a C-terminal TIR domain (Fig. S1A). The DD is essential for the recruitment of interleukin-1 receptor-associated kinase (IRAK) family members, particularly IRAK4, IRAK1 and IRAK2.^8,10,11^ These IRAK kinases feature an N-terminal DD and a C-terminal serine/threonine kinase or pseudokinase domain.^10,11^ Previous structural studies have elucidated the arrangement of the MyD88^DD^−IRAK4^DD^−IRAK2^DD^ complex revealing a helical structure termed the ‘Myddosome’ with a stoichiometry of 6:4:4.^12^ This molecular arrangement brings the kinase domains into close proximity promoting their transphosphorylation and activation.^13^ Once activated, these kinases recruit TRAF6 to the Myddosome, ultimately leading to activation of the nuclear factor κ-light-chain-enhancer of activated B cells (NF-κB) and the MAP kinase pathways and induction of a pro-inflammatory response.^8,9,14,15^

Patients with autosomal recessive deficiency of the *MyD88* gene due to deletion or missense mutations in both chromosomes are highly susceptible to pyogenic bacterial infections which can have an early onset in childhood.^16^ By contrast somatic gain-of-function mutations have been detected in patients suffering from lymphomas. Notably, the L252P mutation (historically referred to as L265P) is the most frequent somatic mutation in lymphoid cancers, found in approximately 10-20% of patients with activated B-cell-like diffuse large B-cell lymphoma (ABC-DLBCL) and in 90% of patients diagnosed with Waldenström macroglobulinemia (WM).^16–19^ The L252P mutation occurs in a β-sheet in the core of the TIR domain leading to spontaneous punctum formation, recruitment and activation of IRAK4, IRAK1 and IRAK2, and downstream pathway activation, ultimately promoting cell survival.^17,20^ Understanding the effect of this oncogenic driver mutation, L252P, may be a good starting point for developing better therapeutics and improving patient outcomes in the future.

Several monomeric structures of MyD88^TIR^ have been solved by nuclear magnetic resonance (NMR) (PDB 2Z5V and 2JS7).^21^ In addition, structures of MyD88^TIR^ by X-ray crystallography (PDB 4DOM and 4EO7)^22^ and microcrystal electron diffraction (MicroED) (PDB 7BEQ and 7L6W)^23^ revealed linear filament structures of MyD88^TIR^ in the crystal packing; these filament structures are similar to the basic TIR filament assembly observed in Mal^24^, AbTir^25^ (TIR domain-containing protein from *Acinetobacter baumannii*) and SPARTA^26^ [short prokaryotic argonaute TIR-APAZ (analogue of Piwi–Argonaute-Zwille domain)]. However, it remains unclear whether these MyD88 filament structures observed from crystal packing represent the activated and signaling competent form of MyD88. Moreover, the structure of the oncogenic L252P mutant MyD88 has not yet been determined.

To address these knowledge gaps, we determined the cryo-EM structures of the wild-type (WT) and cancer-specific L252P mutant MyD88 TIR domains at resolutions of 3.6 Å and 3.4 Å, respectively. Our findings reveal that both WT and L252P TIR domains form double-filament assemblies arranged in an antiparallel fashion, but with WT organized into rings and L252P into spirals. Unexpectedly, the MyD88^TIR^ assemblies are bound with double-stranded DNA (dsDNA). Combined with biochemical assays, cellular experiments, and RNA sequencing (RNA-seq), we revealed an unexpected DNA-binding activity of MyD88 and investigated its contribution to cytosolic DNA signaling.

### MyD88^TIR^ forms double-filament rings

To gain a deeper understanding of MyD88^TIR^ assembly, we sought to solve its structure using cryo-EM. We tested various construct boundaries and ultimately expressed the human MyD88^TIR^ domain (residues 155-296) fused N-terminally to an His-MBP-TEV tag in bacteria (Figure S1B). Gel filtration chromatography revealed a fraction of the protein eluted at the void volume with an elevated A_260_/A_280_ ratio (Figure S1C,D). Using negative-stain EM, we observed ring-like structures with an average diameter of approximately 30 nm (Figure S1E).

Using this sample from the void peak, we prepared cryo-EM grids and collected a large dataset (Figure S1F-I). Because these rings are heterogeneous in size and shape, we picked segments of the rings for structure determination, like what we did for the structure determination of NINJ1 oligomers^27^. Two-dimensional (2D) classification of the ring segments revealed a repetitive pattern within the ring, with a fuzzy density on the convex side corresponding to the MBP tag and a thin density on the concave side (Figure S1G). An initial three-dimensional (3D) reconstruction revealed a double-ring assembly, and refinement with C2 symmetry resulted in a structure of this ring segment at a resolution of 3.6 Å (Figures 1A and S1I,J, Table S1 and Video S1).

**Figure 1.**
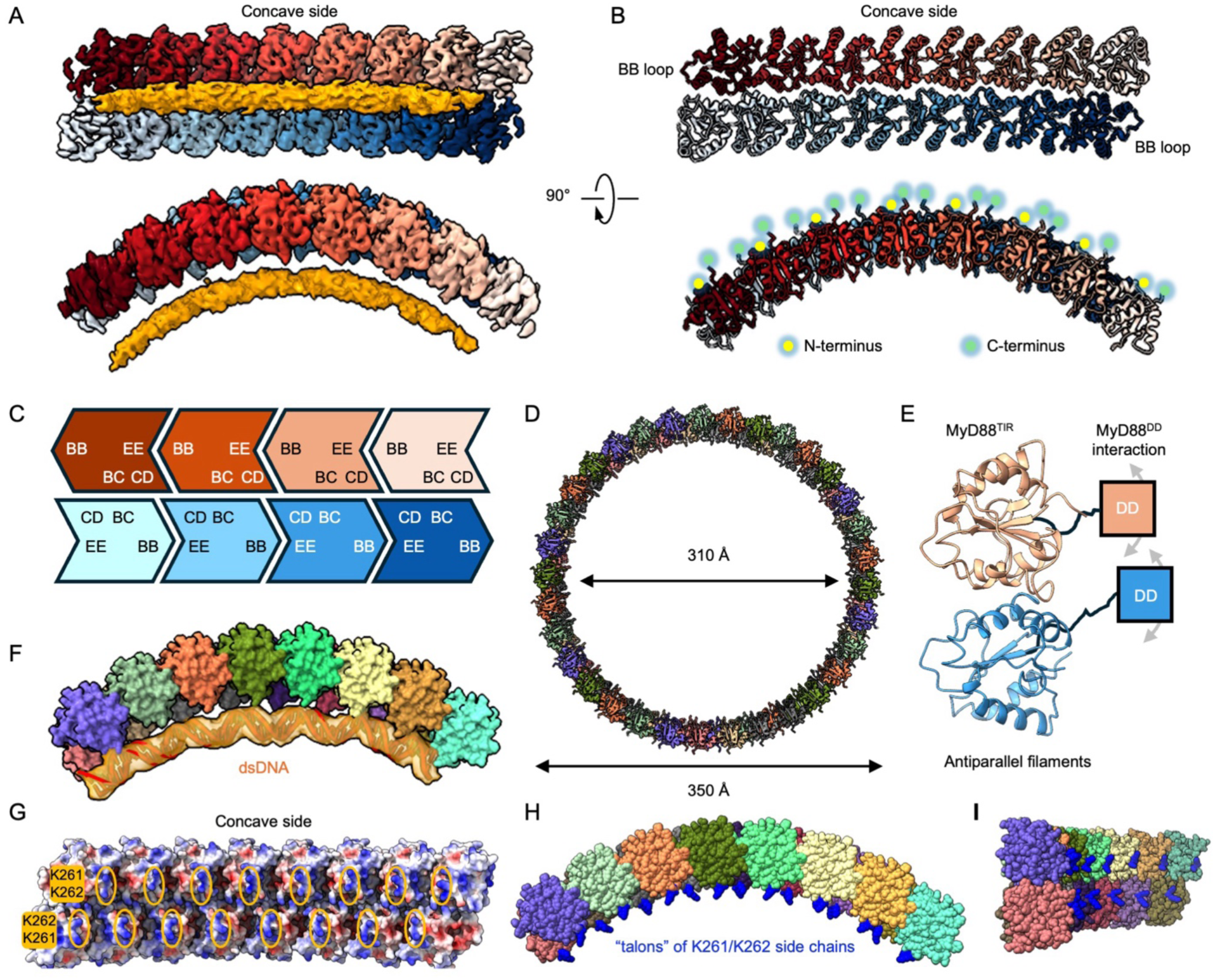
Structure of the MyD88^TIR^ ring bound to dsDNA. **(A)** Cryo-EM map of the 16mer MyD88^TIR^ double-layer ring segment in two orientations and with individual TIR domains coloured in different shades of red and blue, respectively. An additional density is shown in orange. **(B)** Atomic model of the antiparallel MyD88^TIR^ ring segment in two orientations and colour-matched to (A), with N- and C-termini extending outward to the convex ring side. **(C)** Schematic of an octameric MyD88^TIR^ ring segment showing the interaction elements at the interfaces. **(D)** Model of a MyD88^TIR^ ring comprised of 66 subunits and constructed from the 16mers. The inner and outer dimensions are approximately 310 and 350 Å, respectively. **(E)** MyD88 death domains (DDs) from all subunits reside at the same side of the antiparallel double-layered segment. **(F)** Cryo-EM map from 3D classification with a focused mask (yellow) around the extra density at the concave side; a fitted HSV-60bp dsDNA model (orange) is shown within the transparent mask density. **(G)** Electrostatic surface potential of the TIR ring concave side that faces dsDNA, with K261 and K262 residues highlighted. **(H)** Top view of the 16mer with each subunit in a different colour and K261 and K262 in blue, showing the talon-like side chains of the lysine residues for gripping dsDNA. **(I)** A slanted side view of the 16mer in (H).

We built the nearly complete atomic model of MyD88^TIR^ spanning residues 161-296, revealing an arc structure of 16 subunits with two curved filaments running antiparallel to each other and a C2 symmetry axis perpendicular to the arc (Figure 1B,C). Sequential alignment of the hexadecameric model reveals a fully closed double-layered ring with each layer composed of 33 MyD88^TIR^ protomers and inner and outer diameters of 310 Å and 350 Å, respectively (Figure 1D). The TIR domain conformation is highly conserved with a Cα root-mean square deviation (RMSD) of 0.8 Å over 129 Cα atoms compared to the previously determined MyD88 MicroED crystal structure^23^ (PDB ID: 7BEQ) (Figure S2A). However, in the crystal lattice of the latter, the packing-inferred straight filaments are arranged in a parallel, rather than antiparallel orientation, whereas TIR structures from other mammalian or lower species can adopt either arrangement^23,24^ (Figure S2B). The antiparallel filament orientation in our MyD88 ring positions the N-termini of both rings where the MyD88 DDs are located on the same convex side to promote DD oligomerization and recruitment of downstream signaling kinases IRAK4 and IRAK2 (Figures 1E and S2C).

Within each filament of the ring, interactions between adjacent subunits occur via interaction of the BB loop (the loop connecting βB and αB) of one subunit with the EE surface (consisting of the βD and βE strands and the αE helix) (Figure 1C), which are consistent with those reported previously in the MyD88^TIR^ filaments observed in crystal packing.^23^ The interstrand interactions in the MyD88 ring occur predominantly via the BC and CD interfaces consisting of the αB and αC helix as well as the CD loop (Figures 1C and S4). By contrast, the antiparallel TIR domain assembly of human SARM1 (sterile alpha and Toll/interleukin-1 receptor motif-containing 1) engages interstrand interactions at different surfaces (Figure S2B).

The interstrand interactions in our MyD88^TIR^ structure are mainly hydrophobic interactions involving residues W205 of the αB helix, F235 and F239 of the αC helix, and I267 and F270 of the αD helix (Figure S3). Moreover, the aliphatic side chains of K231 and K238 engage the neighbouring subunit and are part of the interstrand interface (Figure S3). Interestingly, the S242 side chain projects into a partially hydrophobic pocket formed by F235 and F239 of the neighbouring subunit (Figure S3). S242 has been reported as an inhibitory phosphorylation site for downstream NF-κB activation.^20,28^ Based on our structure, introducing a large, negatively charged moiety at this position would create steric clashes and unfavourable interactions with the neighbouring subunit, likely destabilizing double-layered ring formation.

### The MyD88^TIR^ ring is bound with dsDNA

After completion of the model building, we could not assign any residues from either the MyD88^TIR^ or the purification tags to the thin density on the concave side of the ring assembly (Figure 1A). To further improve the resolution of this unidentified density, we conducted focused 3D classification using a soft mask covering this extra density (Figure S4A). The classification revealed considerable variability on this part of the structure, both in terms of its appearance and occupancy (Video S2). Despite this heterogeneity, we successfully isolated one class that exhibited an improved density with more distinct features (Figure S4A). Upon closer examination, the shape and dimensions of this density strongly resembled a helical shape that is consistent with the characteristic dimensions of double-stranded DNA (Figure S4B). This finding aligns with the high A_260_/A_280_ absorbance ratio we observed in gel filtration chromatography (Figure S1C), which is indicative of the presence of nucleic acids. We also loaded and separated an aliquot used to prepare cryo-EM grids on an agarose gel and visualized a smear of nucleic acids; if they were dsDNA species, they mainly represented sizes of around ∼750 bp and larger, likely genomic fragments at the top of the gel (Figure S4C). Furthermore, our cryo-EM micrographs revealed the presence of thin filamentous structures that shared similar dimensions to the additional density observed in our reconstructions (Figures S4D,E). We manually picked these thin filamentous particles and solved their structure at ∼6 Å resolution (Figure S4F). The resulting map allowed us to unambiguously fit a model of double-stranded DNA, with clear evidence of the expected helical pitch (Figure S4F).

We fitted a 60 bp dsDNA model into the additional thin density observed in our MyD88^TIR^ reconstruction (Figure 1F), which aligns along the inner surface of the double-layered TIR ring. The concave surface of the antiparallel MyD88^TIR^ ring features positively charged patches primarily contributed by two lysine residues from each subunit, K261 and K262 (Figure 1G), whose side chains protrude like talons to engage and grip the dsDNA phosphate backbone (Figure 1H,I). Comparative analysis of surface charge distributions among human TIR domains revealed that MyD88 exhibits a uniquely positive surface charge, unlike other mammalian TIR domains that typically display negatively charged surfaces or concentrate positive charges at TIR–TIR interaction sites (Figure S5). Based on the location of the dsDNA density, the lysine side chains appear positioned to interact exclusively with the phosphate backbone (Figure S6A). Thirty-two lysine residues could engage approximately 60 bp of dsDNA, corresponding to a spacing of ∼28 Å per four lysines. In the double-layered ring, regularly spaced positively charged patches along the filament lengths grip dsDNA from two sides. These intriguing data suggest the possibility of a specialized role of MyD88 in DNA recognition not shared by other TIR domains.

A closer inspection of our micrographs revealed extended stretches of dsDNA protruding from the lateral edges of the MyD88^TIR^ rings (Figure S4D), suggesting that DNA sequences exceeding ∼240 bp cannot be fully enclosed within a single ring. This observation implies that while each MyD88^TIR^ ring can accommodate ∼240 bp of dsDNA, any additional length could extend beyond the ring structure from either end (Figure S4E). Such a configuration may be particularly relevant during bacterial or viral infections, where longer DNA fragments are common. Moreover, it is tempting to speculate that multiple MyD88^TIR^ rings could assemble along a single dsDNA molecule, potentially modulating the immune response in a length-dependent manner.

### MyD88^TIR^ binds dsDNA *in vitro*

To gain deeper insights into the MyD88–dsDNA interaction, we performed electrophoretic mobility shift assays (EMSAs) to assess the binding affinities using dsDNA of varying lengths. MBP-MyD88^TIR^, purified from the later, presumably monomeric size-exclusion fractions, was incubated at 37 °C with Cy3-labelled dsDNA fragments of 30, 45, 60, and 75 base pairs (bp) derived from Herpes Simplex Virus-1 (HSV-1) sequences, and the complexes were resolved by native PAGE. MBP-MyD88^TIR^ shifted dsDNA in a concentration-dependent manner (Figures 2A and S6B). The binding affinities for the different DNA lengths were broadly similar, all showing evidence of cooperativity as indicated by the fitted Hill coefficients (Figures 2B,C). Competition EMSAs using unlabeled DNA further confirmed the specificity of the interaction and excluded non-specific binding to the Cy3 fluorophore (Figures 2D,E and S6C).

**Figure 2.**
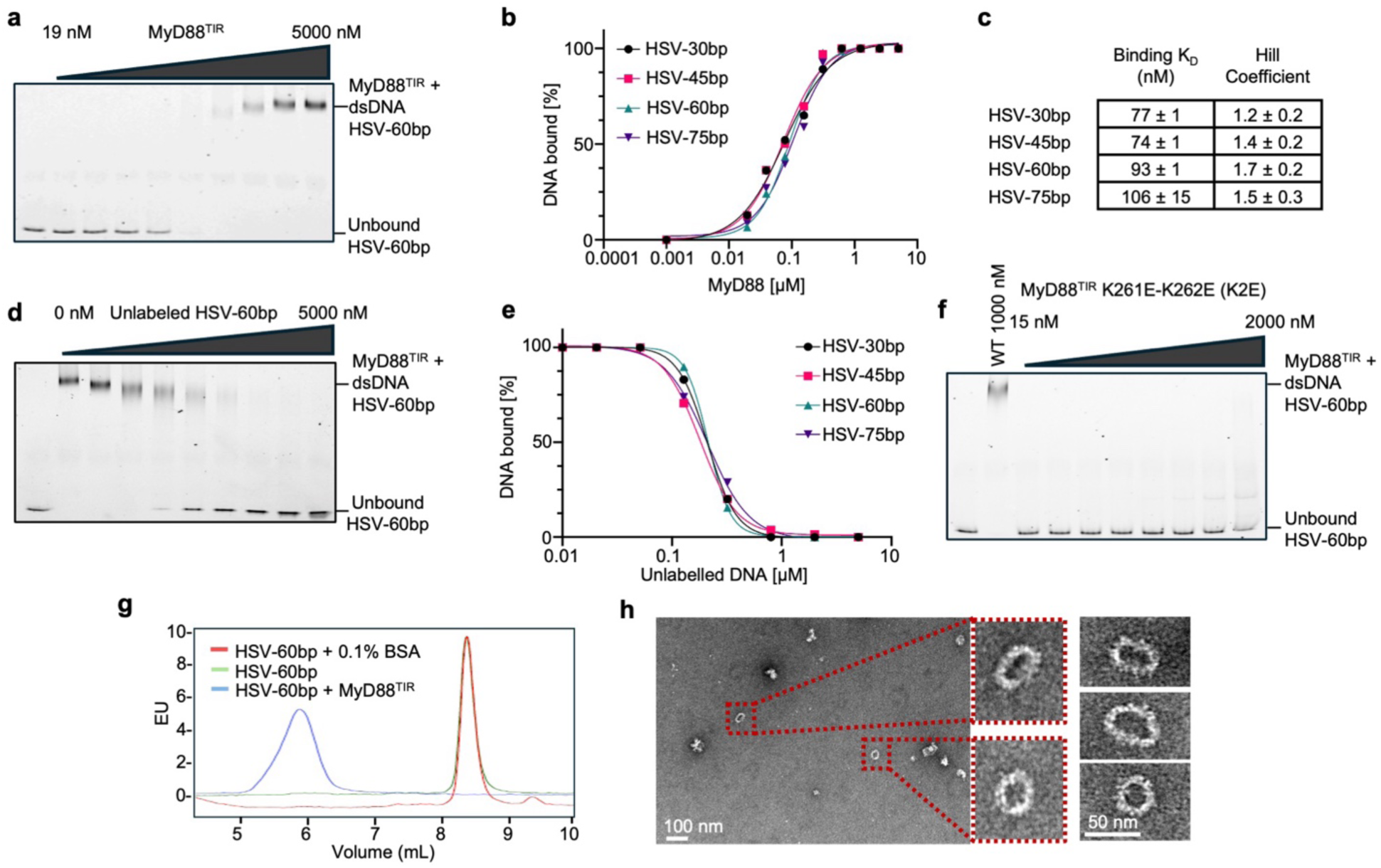
Electrophoretic mobility shift assays. **(A)** EMSA of Cy3-labeled HSV-60bp dsDNA (0.5 nM) with increasing concentrations of MyD88^TIR^ (19, 39, 78, 156, 312, 625, 1,250, 2,500 and 5,000 nM). **(B)** Quantification of DNA bound from (A) and other lengths of HSV-1 dsDNA (30, 45, and 75 bp). **(C)** Binding KD and Hill coefficient derived from EMSAs (A, B). **(D)** Competition with increasing amount of unlabelled HSV-60bp dsDNA (0, 8, 20, 51, 128, 320, 800, 2,000 and 5,000 nM) in EMSA of MyD88TIR (2,000 nM) and Cy3-labeled HSV-60bp dsDNA (0.5 nM). **(E)** Quantification of DNA bound from (D) and other lengths of HSV dsDNA (30, 45, and 75 bp). **(F)** EMSA of Cy3-labeled HSV-60bp dsDNA (0.5 nM) with increasing amount of the MyD88^TIR^ K2E mutant (15, 31, 62, 125, 250, 500, 1,000 and 2,000 nM), showing no apparent shift. MyD88^TIR^ (1,000 nM) was used as a positive control. **(G)** Fluorescence-detection size-exclusion chromatography of Cy3-labeled HSV-60bp dsDNA. Monomeric His-MyD88^TIR^ (20 nM) or control BSA (0.1%, or 15 µM) was incubated at 37 °C with 20 nM Cy3-labeled HSV-60bp dsDNA for 1 h. Reaction buffer contains 40 mM Hepes at pH 7.0 and 150 mM NaCl. **(H)** Negative-stain EM image of full-length His-tagged MyD88 sample from the void volume.

To determine whether dsDNA binding is mediated by K261 and K262, as suggested by the structure, we introduced charge-reversal mutations in MBP-MyD88^TIR^ (K261E/K262E; hereafter referred to as K2E). The K2E mutation completely abrogated the MyD88–dsDNA interaction in the EMSA (Figure 2F), supporting the structural model of dsDNA engagement. Additionally, to rule out any influence of the MBP tag on the interaction, we performed fluorescence size-exclusion chromatography (FSEC) using His-tagged recombinant MyD88^TIR^ and Cy3-labelled HSV-60bp dsDNA. In the presence of MyD88^TIR^, Cy3-HSV-60bp dsDNA eluted in earlier fractions, confirming the formation of a complex between MyD88 and dsDNA (Figure 2G). Finally, recombinant full-length His-tagged MyD88, despite low yield, formed similar ring structures under negative-stain EM (Figure 2H) and retained the ability to interact with IRAK4 (Figure S6D), supporting the functionality of MyD88 rings in absence of a large affinity tag.

### Cytosolic dsDNA activates MyD88-dependent cytokine expression

MyD88 is primarily known for its role as an adaptor protein for TLR- and IL-1R-dependent inflammatory gene expression.^4^ These pathways involve activation of IRAK family kinases, which stimulates the E3 (ubiquitin ligase) activity of TRAF6, leading to ubiquitin-dependent activation of the NF-κB pathway.^29^ Given the direct MyD88–dsDNA interaction, we asked whether dsDNA could trigger MyD88 activation to initiate NF-κB transcriptional regulation. To investigate these questions, we first examined the mRNA and the secreted protein levels of TNFα, a classical inflammatory gene.^30,31^ We found that MyD88 is required to increase *Tnf* mRNA level and TNFα protein secretion in mouse immortalized bone marrow-derived macrophages (iBMDMs) upon transfection of HSV-60bp dsDNA (Figures 3A,B). By contrast, knockout (KO) of cGAS, the known cytosolic DNA sensor for activating type-I interferon gene^32,33^, minimally affected TNFα expression (< 2-fold), and pretreatment with the potent cGAS inhibitor G140^34^ in MyD88 KO iBMDMs had no additional effects (Figures 3A,B and S7A). When we increased the dsDNA lengths to 2 kb and 7 kb, TNFα expression was further increased in comparison with HSV-60bp dsDNA, which was however equally dependent on cGAS and MyD88 (Figure S7B). Thus, dsDNA-induced expression of the inflammatory TNFα depends on both cGAS and MyD88.

**Figure 3.**
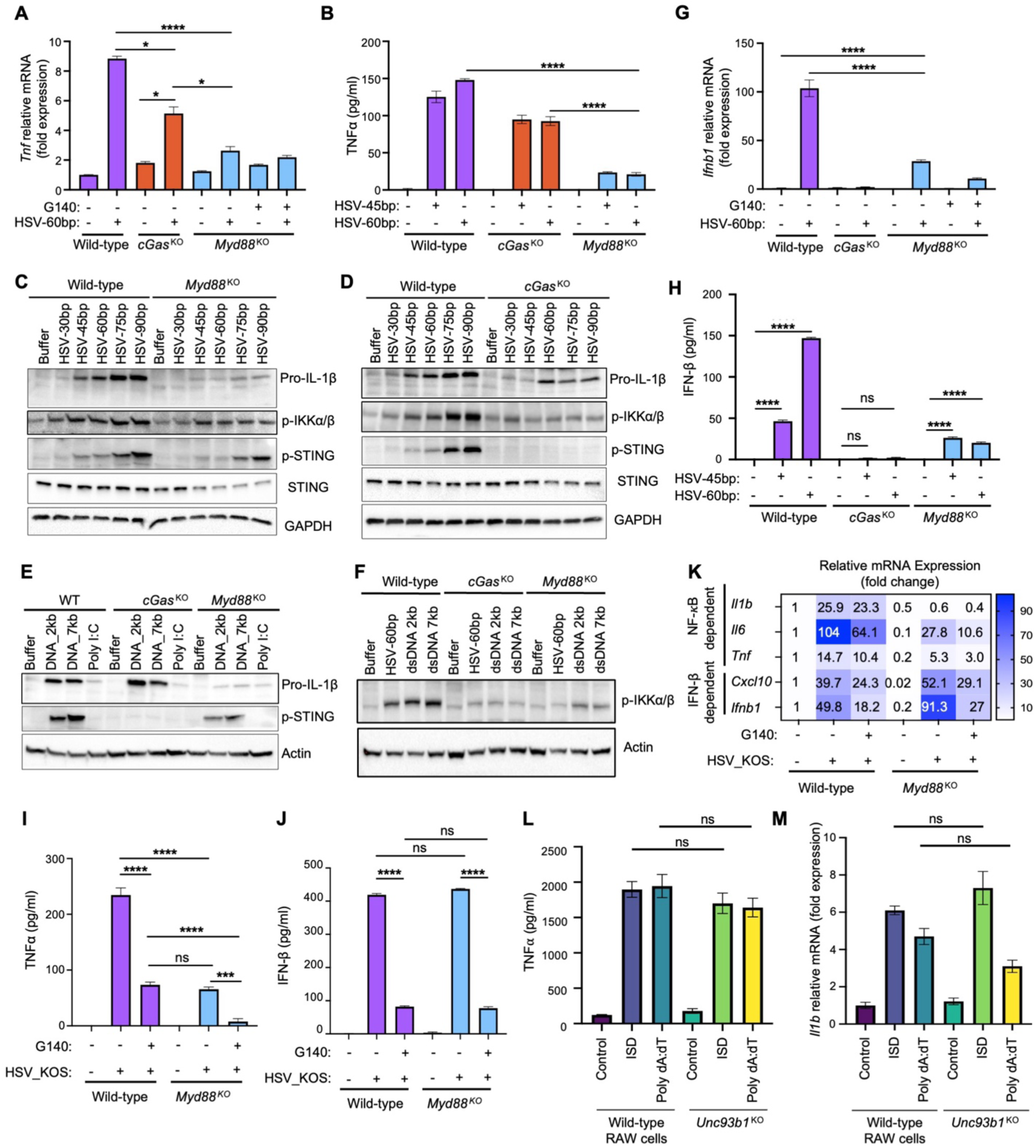
MyD88 is a sensor for dsDNA-mediated NF-κB activation. **(A)** Fold induction of *Tnf* mRNA expression analysed by qRT-PCR. WT, *Myd88* KO, or *cGas* KO iBMDMs were transfected with HSV-60bp dsDNA (2.5 µg) for 6 h. The indicated *Myd88* KO iBMDMs were pretreated with the cGAS inhibitor G140 (10 µM) for 12 h. **(B)** TNFα release analysed by ELISA. WT, *Myd88* KO, or *cGas* KO iBMDMs were transfected with HSV-45bp or HSV-60bp dsDNA (2.5 µg) for 6 h. (**C, D)** Pro-IL-1β, phospho-IKKα/β, phospho-STING and total STING levels analysed by western blots. WT and *Myd88* KO (C) or WT and *cGas* KO iBMDMs (D) were transfected with increasing lengths of HSV-1 dsDNA (30 to 90 bp) (2.5 µg) for 6 h. **(E)** Pro-IL-1β and phospho-STING levels analysed by western blots. WT, *cGas* KO, or *Myd88* KO iBMDMs were transfected with poly I:C (2.5 µg) or two dsDNA fragments (2.5 µg) corresponding the length boundaries of poly I:C, 2,000 bp and 7,000 bp respectively, for 6 h. **(F)** Phospho-IKKα/β levels analysed by western blots. WT, *cGas* KO, or *Myd88* KO iBMDMs were transfected with HSV-60bp dsDNA, 2,000 bp or 7,000 bp dsDNA, for 6 h. **(G)** Fold induction of *Ifnb1* mRNA expression analysed by qRT-PCR. WT, *Myd88* KO, or *cGas* KO iBMDMs were transfected with HSV-60bp dsDNA (2.5 µg) for 6 h. The indicated *Myd88* KO cells were pretreated with G140 (10 µM) for 12 h. **(H)** IFN-β release analysed by ELISA. WT, *cGas* KO or *Myd88* KO iBMDMs were transfected with HSV-45bp or HSV-60bp dsDNA (2.5 µg) for 6 h. **(I, J)** TNFα (I) or IFN-β (J) release analysed by ELISA. WT or *Myd88* KO primary BMDMs were infected with HSV-1 KOS strain for 18 h. The indicated cells were pretreated with G140 (10 µM) for 12 h. (**K**), Relative mRNA fold change of selective NF-κB and IFN-β dependent genes measured by qRT-PCR and plotted. Infection condition is the same as in (I, J). **(L, M)** TNF*a* secretion determined by ELISA (L) and fold induction of *Il1b* analysed by qRT-PCR (M). WT or *Unc93b1* KO RAW 264.7 macrophages were transfected with inflammatory stimulatory DNA (ISD, 2.5 µg) or poly dA:dT (2.5 µg) for 6 h.

We next tested dsDNA-induced pro-IL-1β expression. Transfection of HSV-1 dsDNA caused pro-IL-1β expression in a length-dependent manner from 30bp to 90bp (Figure 3C). MyD88 KO almost abolished this dsDNA-triggered pro-IL-1β expression (Figure 3C). cGAS KO diminished but did not abolish pro-IL-1β expression (Figure 3D). When the dsDNA lengths were increased to 2 and 7 kilo bp (kb), MyD88 KO abolished, while cGAS KO had no effect on, the dsDNA-triggered pro-IL-1β expression (Figure 3E). Both MyD88 KO and cGAS KO dampened IKKα/IKKβ phosphorylation upon DNA transfection (Figures 3C,D,F). Transfection with poly I:C, dsRNA of similar lengths (1.5 kb - 8 kb), did not activate expression of the NF-κB-dependent gene pro-IL-1β in WT, MyD88 KO, or cGAS KO iBMDMs (Figure 3E), suggesting that MyD88 does not sense dsRNA.

As controls, we tested IFN-β gene expression by mRNA and secretion levels in MyD88 KO or cGAS KO iBMDMs. As expected^35^, IFN-β mRNA and protein levels were completely abolished by cGAS KO upon transfection of two different lengths of HSV-1 dsDNA or kb long dsDNA (Figures 3G,H and S7C). Surprisingly, MyD88 KO also significantly dampened the levels of *Ifnb1* mRNA and IFN-β secretion, and G140 pre-treatment further lowered *Ifnb1* mRNA (Figures 3G,H and S7C). Consistent with a previously implicated role of MyD88 in inhibiting p-STING degradation, total STING and p-STING levels in MyD88 KO iBMDMs were both reduced upon transfection of HSV-1 dsDNA (Figure 3C). By contrast, in cGAS KO iBMDMs, there was abolished p-STING but no observable total STING reduction (Figure 3D). Together with the RNA-seq data (see below), our findings suggest that the reduced IFN-β levels in MyD88 KO cells may result from dual roles of MyD88 in promoting *Ifnb1* transcription and preventing p-STING degradation.

### Inflammatory cytokine expression by HSV-1 infection is dependent on MyD88

To examine our finding in a more physiological setting, we carried out HSV-1 infection of primary BMDMs obtained from WT and MyD88 KO mice and analysed TNFα and IFN-β expression. HSV-1 infection induced TNFα expression in primary WT BMDMs, whereas this response was significantly reduced in MyD88 KO BMDMs, highlighting the importance of MyD88 in mediating the inflammatory cytokine response (Figure 3I). Inhibition of cGAS by the small molecule G140 further reduced HSV-1-induced TNFα expression in MyD88-deficient BMDMs, suggesting that cGAS and MyD88 function in parallel or converging pathways to promote TNFα expression upon HSV-1 infection (Figure 3I). By contrast, IFN-β expression by HSV-1 infection of BMDMs was not affected by the absence of MyD88 (Figure 3J), unlike transfection of short HSV-1 dsDNA in iBMDMs (Figure 3H). To further dissect the differential regulation of IFN-β versus NF-κB-dependent gene expression in response to HSV-1 infection, we examined well-established target genes of each pathway. MyD88 deficiency markedly impaired the transcription of NF-κB-dependent proinflammatory cytokine genes (*Il1b*, *Il6*, and *Tnf*), whereas the expression of IRF3-dependent genes such as *Cxcl10* and *Ifnb* remained unaffected (Figure 3K).

We also used the HSV-1 strain 7134, which lacks ICP0, a protein known to downregulate inflammatory response.^36^ This strain can induce cytokine responses at a lower multiplicity of infection (MOI) and has been used to examine the role of cGAS–cGAMP signaling in antiviral defense.^35^ With HSV-1 7134 at an MOI of 10 in primary WT and *Myd88* KO BMDMs, TNFα and IFN-β were both robustly elevated to higher levels than with WT HSV-1 at MOI of 100 (Figures S7D,E and 3I,J). While TNFα activation was highly dependent on MyD88 but not cGAS (Figure S7D), IFN-β activation was independent of MyD88 (Figure S7E). These data supported that HSV-1-induced inflammatory cytokine expression, but not interferon expression, is highly dependent on MyD88.

### *Unc93b1* deficiency does not dampen inflammatory cytokine expression

Because TLR9, which recognizes ssDNA entities with motifs containing the unmethylated CG sequence (CpG), uses MyD88 for its signaling^37^, we tested if TLR9 was responsible for the MyD88-dependent cytokine expression upon dsDNA transfection. We first used the TLR9 inhibitor ODN2088. While ODN2088 inhibited activation of TLR9 by the agonist ODN1826 (Figure S7F), it failed to inhibit dsDNA-induced MyD88 activation (Figure S7G). It is known that endosomal nucleic acid-sensing Toll-like receptors (TLRs), specifically TLR3 (dsRNA), TLR7-8 (ssRNA), and TLR9 rely on the membrane protein UNC93B1 for their trafficking, stability, and signaling.^38–40^ Because of the availability of *Unc93b1* KO RAW 264.7 monocytes/macrophages^41^ (Figure S7H), we tested if dsDNA-induced inflammatory cytokine expression is dependent on UNC93B1. Production of TNF*a* and *Il1b* upon dsDNA transfection showed no difference in control versus *Unc93b1* KO RAW 264.7 cells (Figures 3L,M and S7I). As a control, we also analysed *Ifnb1* and *Cxcl10*, two genes that are cGAS-dependent, and showed lack of change by *Unc93b1* deficiency (Figure S7I). These results supported that dsDNA transfection-induced inflammatory cytokine production is not dependent on nucleic acid-sensing TLRs such as TLR9.

### DNA binding is required for MyD88’s response to cytosolic DNA but not in TLR signaling

To investigate the direct interaction between cytosolic DNA and MyD88 in cells, we transfected dsDNA into the previously generated iBMDMs expressing GFP-tagged endogenous MyD88.^31^ DNA transfection induced the formation of punctate MyD88 structures, which were not observed in buffer-transfected control cells (Figure 4A). In agreement with our findings, MyD88 punctum structures colocalized with dsDNA (Figure 4A), implicating a direct role of MyD88 in sensing and responding to cytosolic DNA. We further used stimulated emission depletion microscopy (STED) to observe in more detail the micron-sized puncta in these cells. Similar to Myddosome puncta formed by TLR activation^31^, dsDNA-induced Myddosome formation involved barrel-shaped distribution of both DNA and MyD88 in these puncta (Figure S8A).

**Figure 4.**
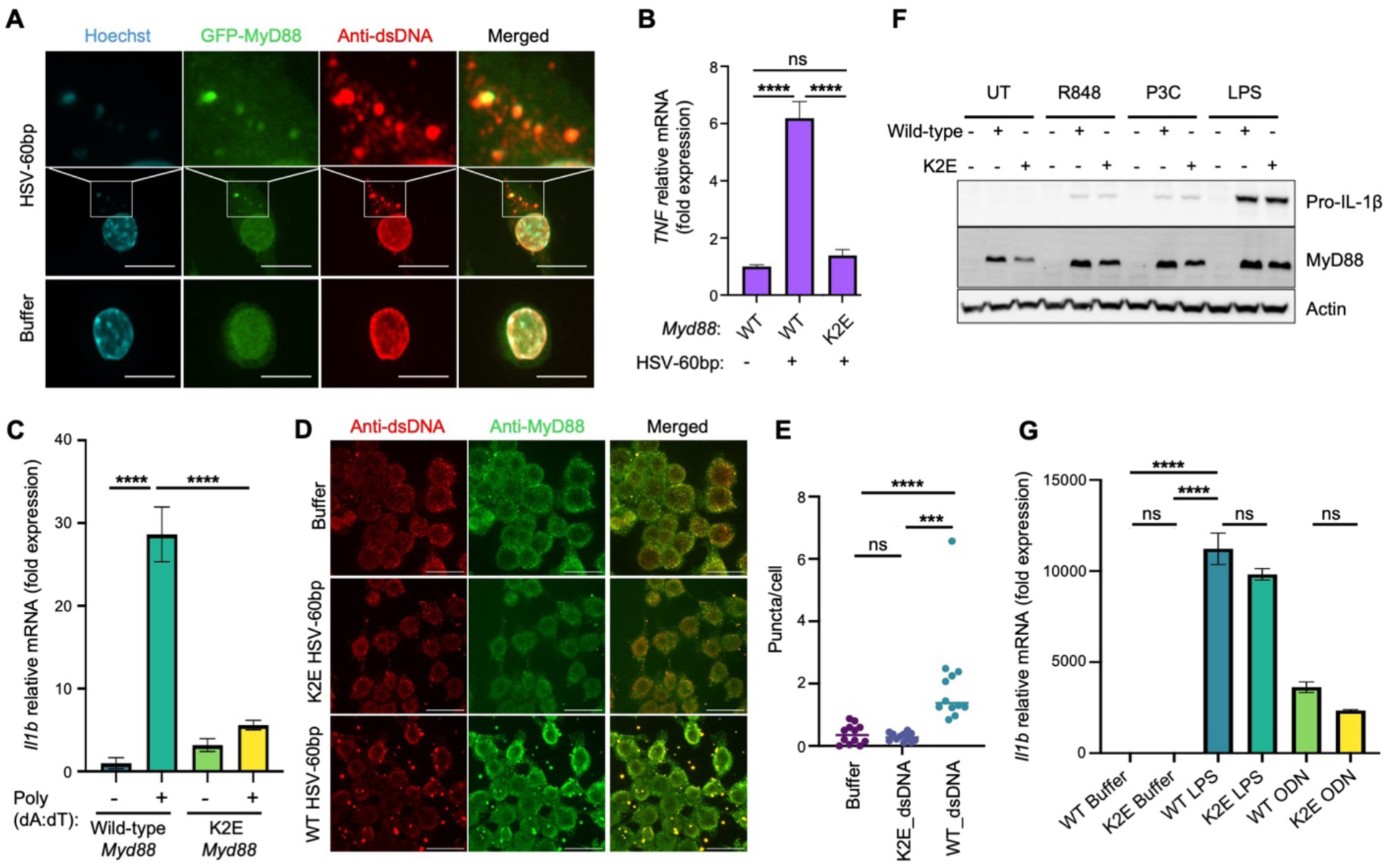
DNA binding mutation reveals binding is required for MyD88’s response to cytosolic DNA. **(A)** Colocalization of MyD88 and dsDNA. WT iBMDMs with endogenously GFP- and FLAG-tagged MyD88 were transfected with HSV-60bp dsDNA for 6 h. The cells were fixed and visualized with GFP fluorescence and anti-dsDNA immunofluorescence. Scale bar = 10 µm. **(B)** *Tnf* mRNA fold change analysed by qRT-PCR. WT *Myd88* KO, or K2E *Myd88* iBMDMs were transfected with HSV-60bp dsDNA (2.5 µg) for 6 h. **(C)** Fold induction of *Il1b* mRNA expression analysed by qRT-PCR. *Myd88* KO iBMDMs reconstituted with WT or K2E Myd88 were transfected by poly dA:dT dsDNA for 6 h. **(D)** Impairment of co-localization with dsDNA in the K2E mutant. *Myd88* KO iBMDMs reconstituted with WT or K2E *Myd88* were transfected with HSV-60bp dsDNA for 6 h. The cells were fixed and stained with anti-MyD88 and anti-dsDNA antibodies and imaged by confocal microscopy. Scale bar = 20 µm. **(E)** MyD88 puncta in confocal images (D) were detected after Gaussian blur (σ = 4.0), quantified, and normalized to the total number of cells per field of view for each condition. **(F)** *Myd88* KO iBMDMs reconstituted with WT or K2E *Myd88* were stimulated by various TLR agonists: R848 for TLR7/8, P3C for TLR2/TLR1 heterodimer, and LPS for TLR4. UT: untreated control. Pro-IL-1β and MyD88 levels were analysed by western blots. Data in bar graphs are represented as mean ± SEM. **(G)** *Myd88* KO iBMDMs reconstituted with WT or K2E *Myd88* were stimulated by the TLR4 ligand LPS and TLR9 ligand ODN1826. Fold induction of *Il1b* mRNA expression was analysed by qRT-PCR.

To demonstrate that the observed effects from MyD88-dependent cytosolic dsDNA sensing were driven specifically from the MyD88–DNA interaction, we generated *Myd88* KO iBMDMs expressing the MyD88 K2E mutant with impaired DNA binding (Figure 2F and S8B). While WT MyD88 rescued dsDNA-induced *Tnf* and *Il1b* mRNA levels in *Myd88*-deficient iBMDMs, the K2E mutant was unable to initiate cytokine expression in response to cytosolic dsDNA (Figures 4B,C), indicating that direct MyD88–DNA interaction is essential for mediating dsDNA-induced proinflammatory cytokine expression. We also imaged these iBMDMs using anti-dsDNA and anti-MyD88 antibodies. Upon dsDNA transfection, numerous puncta with colocalized dsDNA and MyD88 were observed in WT MyD88-reconstituted cells. By contrast, few puncta were seen in K2E-reconstituted cells after dsDNA transfection, which is visibly similar to buffer-transfected cells (Figure 4D), arguing that WT MyD88, but not K2E, caused DNA to aggregate. Quantification of the images confirmed the markedly reduced punctum formation when the K2E MyD88 was used (Figure 4E). By contrast, we did not observe any significant difference between WT and K2E mutant iBMDMs in pro-IL-1β expression upon TLR activation: Pam3CSK4 for TLR1/2, lipopolysaccharides (LPS) for TLR4, R848 for TLR7/8, and ODN1826 for TLR9 (Figures 4F,G). Thus, dsDNA-binding residues of MyD88 are selectively required for DNA sensing, effectively decoupling TLR- and DNA-induced MyD88 signaling.

To gain more mechanistic insights in the antiparallel orientation of the TIR filaments, we reconstituted *Myd88* KO iBMDMs with MyD88 interstrand interface mutants. As expected, most of the interstrand mutations abolished the MyD88’s ability to trigger dsDNA-induced *Il6* expression (Figure S8C). We further investigated the role of MyD88 in dsDNA-induced cell death. Our results indicated that the release of lactate dehydrogenase (LDH) following transfection with different lengths of HSV-1 dsDNA was unaffected by *Myd88* deficiency, while NLRP3 activation by LPS and nigericin was dependent on MyD88 due to the requirement for LPS priming (Figure S8D). These data argued that MyD88 does not play a role in dsDNA-mediated cell death.

### RNA-seq reveals a broad contribution of MyD88 in cytosolic dsDNA transcriptional response

To dissect which signaling pathways are regulated by MyD88 and to what extent they intersect with the cGAS-STING pathway, we performed bulk RNA sequencing (RNA-seq) on WT, *Myd88* KO, *cGas* KO, and *cGas/Myd88* dKO iBMDMs under buffer, HSV-60bp or poly dA:dT dsDNA transfection (Figure S9A, Table S2, S3). Principal component analysis (PCA) of RNA species showed close clustering of the triplicates (Figure S9B). Evaluation of upregulated differentially expressed genes (DEGs) suggested that like *cGas*, *Myd88* is a major contributor to gene expression in response to dsDNA with 643/1576 genes and 699/1314 genes dampened in *Myd88* KO among the total of upregulated genes in WT by poly dA:dT transfection and HSV-60bp transfection, respectively (Figure 5A). Of the two RNA-seq datasets using poly dA:dT versus HSV-60bp, the shorter HSV-60bp transfection-induced gene expression appeared to be more dependent on MyD88, with more than half (699 genes) of upregulated genes (1314 genes) in WT by DNA transfection dampened in *Myd88* KO (Figure 5A). *cGas*/*Myd88* KO further dampened the dsDNA-induced gene upregulation (Figure 5A). Thus, although *cGas* was necessary for most dsDNA-responsive genes^35^, it is not sufficient for about half of the upregulated genes, as shown by the *Myd88* KO phenotypes.

**Figure 5.**
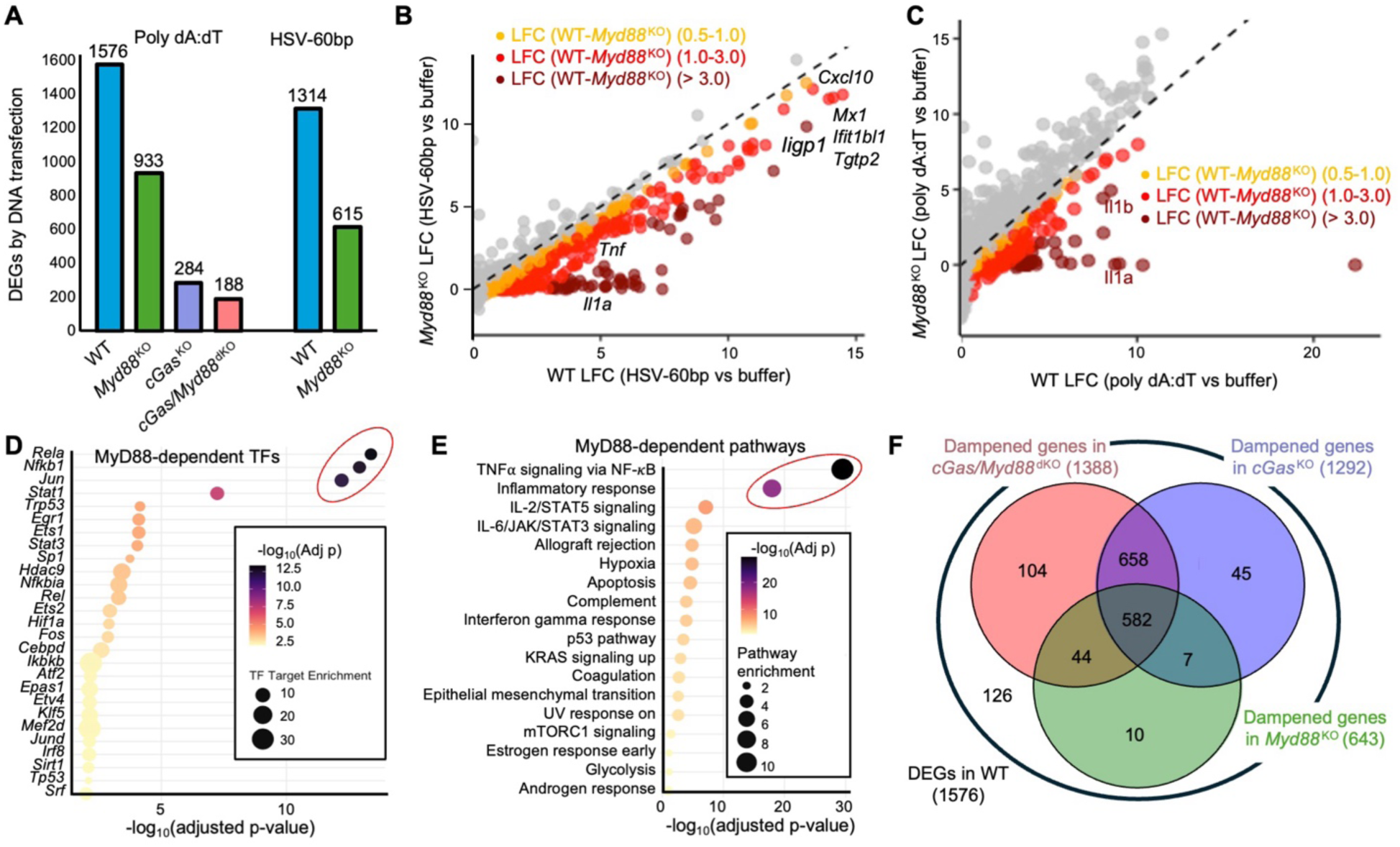
RNA-seq reveals a major role of MyD88 in dsDNA-induced transcriptional activation. **(A)** Number of upregulated differentially expressed genes (DEGs, adjusted p-value < 0.05, |log₂FC| ≥ 0.5) in response to DNA transfection in WT and different genetic KO iBMDMs, revealed by RNA-seq. Two different datasets are labelled with the dsDNA species used. **(B)** A scatter plot of DEGs for log fold change (LFC) in WT (x-axis) and *Myd88* KO (y-axis) iBMDMs upon HSV-60bp transfection. Selected genes are labelled. **(C)** A scatter plot of DEGs for log fold change (LFC) in WT (x-axis) and *Myd88* KO (y-axis) iBMDMs upon poly dA:dT transfection. Selective genes are labelled. **(D, E)** Transcription factor (D) and pathway (E) analyses from genes downregulated by *Myd88* KO. **(F)** Venn diagrams showing the numbers (in parentheses) of downregulated genes in *cGas* KO, *Myd88* KO, and *cGas/Myd88* dKO among the upregulated DEGs in WT iBMDMs upon poly dA:dT transfection.

Plotting of log fold change (LFC) for upregulated DEGs in WT iBMDMs (dsDNA versus buffer transfection) against LFC in *Myd88* KO iBMDMs highlighted genes whose upregulation was dampened by *Myd88* deficiency (Figures 5B,C). Transcription factor analysis of dampened genes in *Myd88* KO revealed overwhelming associations with *RelA, Nfkb1,* and *Jun* (Figures 5D and S9C), which are known transcription factors in the MyD88 pathway. Pathway analysis of the same downregulated genes revealed a dominant enrichment for TNFα signaling via NF-κB and inflammatory response (Figure 5E). For HSV-60bp dsDNA transfection, IFN-*γ* and IFN-*a* induction was also dependent, at least partly, on MyD88 (Figure S9D). A Venn diagram of the knockout data revealed that nearly half of all cGAS-dependent genes are co-dependent on MyD88 (Figure 5F). This subset includes *Il1b* and *Tnf*, which were consistently validated in this study, as well as many additional genes. These results argued for extensive collaboration between cGAS and MyD88 in DNA-sensing induced gene transcription.

### Structure of the cancer specific mutant L252P MyD88^TIR^

Having established a biochemical and structural protocol for the WT MyD88^TIR^ domain, we extended this work by investigating the cancer-specific TIR domain mutant L252P. Gel filtration chromatography showed that majority of the mutant protein eluted broadly over a wide range of elution volumes, suggesting the presence of various oligomeric states (Figure S10A). In parallel, we observed a significant amount of protein eluting at the void volume, indicative of large assemblies (Figures S10B,C). Like for WT MyD88^TIR^, the A_260_/A_280_ absorbance ratio of the void volume fraction was elevated, pointing to the presence of co-purified nucleic acids. Negative-stain EM revealed long, spiral-like filaments for the void fraction sample (Figure S10D), unlike closed rings for WT MyD88^TIR^. We then determined the L252P spiral segment structure at 3.4 Å resolution (Figures 6A,B, S10E-H and Table S1). Like WT MyD88^TIR^, the L252P cryo-EM map contains an additional thin density on the concave side of the spiral segment, which suggests a similar capability of the L252P mutant to bind dsDNA.

**Figure 6.**
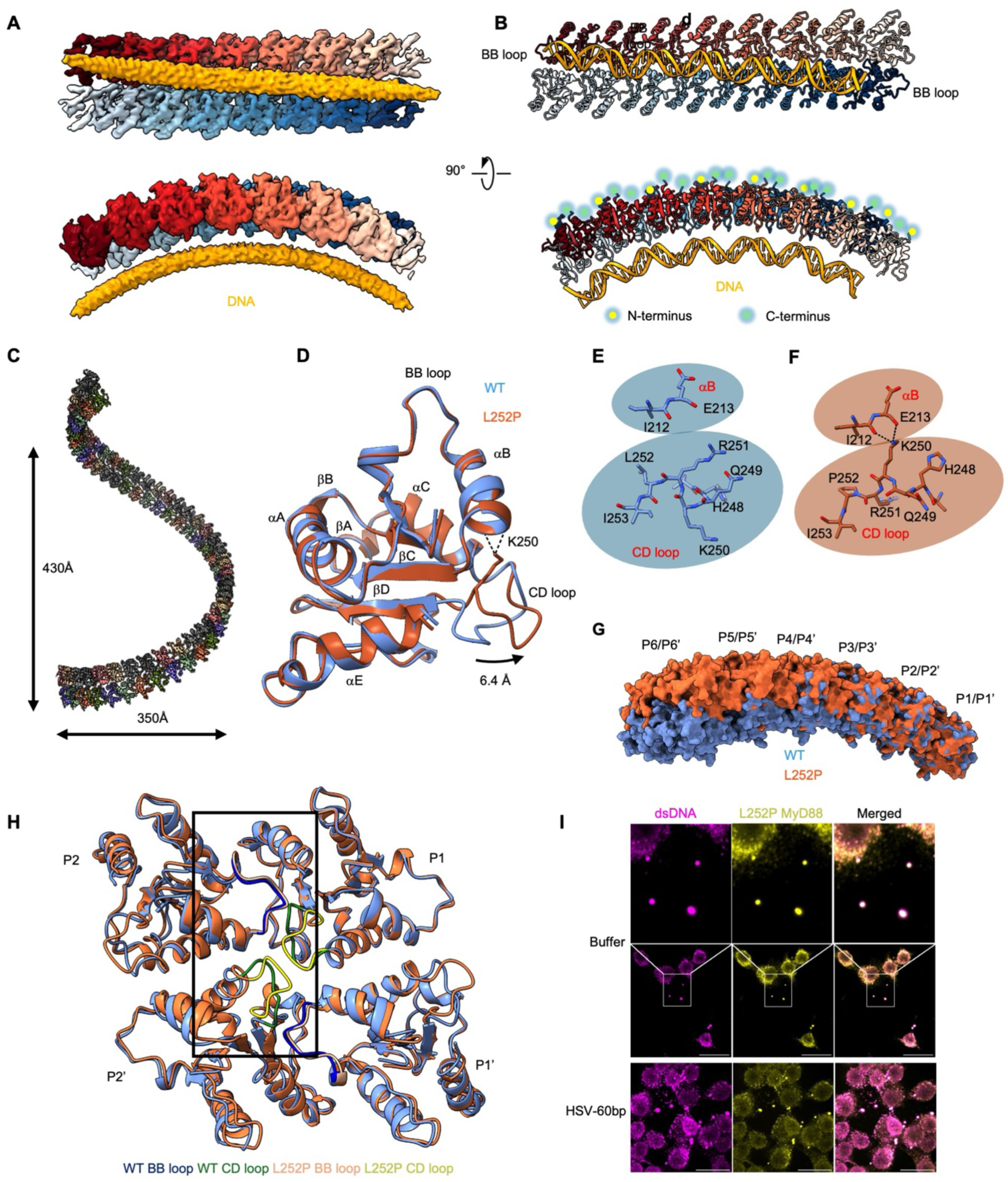
Structure of the MyD88^TIR^ L252P cancer mutant. **A)** Cryo-EM map of the 16mer L252P MyD88^TIR^ ring segment in two orientations and with individual TIR domains coloured in different shades of red and blue, respectively. An additional density is shown in orange. **(B)** Atomic model of the antiparallel L252P MyD88^TIR^ spiral segment in two orientations and colour-matched to (A), with N- and C-termini extending outward to the convex ring side. **(C)** Model of the L252P MyD88^TIR^ spiral with indicated dimensions. **(D)** Overlay of WT (blue) and L252P mutant (orange) MyD88^TIR^ protomers with structural features labelled. The 6.4 Å shift of the CD loop in L252P towards the interstrand interface is indicated. The K250 side chain in L252P is shown to hydrogen bond with the main chain carbonyl oxygens of I212 and E213. (**E, F)** Magnified views of the CD loop in WT (E) and L252P (F) structures, highlighting rearrangements induced by the L252P mutation. **(G)** Surface renderings of WT (blue) and L252P (red) ring segments (12 subunits) overlaid at the left-most subunit of the top layer (P1), showing the gradual deviation of spiral trajectories. P1 and P2 denote subunit positions of the top TIR filament, and P1’ and P2’ denote subunit positions of the lower TIR filament. **(H)** Four WT (blue) and L252P (red) subunits aligned at P1, showing the subtle shifts at P1’, P2 and P2’ subunits induced largely by the CD loop conformation. Within the black box, BB loops are in dark blue and pink for WT and L252P, respectively, and CD loops are in green and yellow for WT and L252P, respectively. **(I)** Colocalization of L252P MyD88 and dsDNA without DNA transfection. *Myd88* KO iBMDMs were reconstituted with L252P *Myd88* and transfected with buffer or HSV-60bp dsDNA for 6 h. Cells were fixed and visualized with anti-MyD88 and anti-dsDNA immunofluorescence. Spontaneous puncta containing L252P and dsDNA were present even with buffer transfection.

The overall oligomeric assembly of the mutant is similar as that of the WT TIR ring. The L252P TIR domain arranges in a double-layered filament with antiparallel orientation of the individual strands. We also placed a model of HSV-60bp dsDNA into the additional density on the concave side of the spiral (Figure 6B), which is the exact same interface used for interaction observed in the WT. When the maps of the L252P mutant are sequentially aligned, a spiral structure is observed with a width of ∼350 Å and a height for one complete turn of ∼430 Å (Figure 6C), highly reminiscent with the appearance on our cryo-EM grids (Figure S10E).

The overall conformation of the WT and L252P MyD88^TIR^ structures are highly similar, with an RMSD of 0.703 Å over 129 Cα atoms, indicating that differences may rest in detail (Figure 6D). A closer look revealed a shift of the CD loop (residues P245–R251) of L252P, which is displaced by approximately 6.4 Å from the TIR domain core of the WT MyD88 towards the interstrand interface (Figure 6D-F and Video S3). Within the CD loop, the shift also causes a 180° flip or switch of the K250 and R251 side chains (Figure 6E,F). The flipped K250 side chain forms new hydrogen bonds with the backbone carbonyl oxygen atoms of I212 and E213 in the critical αB helix (Figure 6D,F). We propose that K250-mediated hydrogen bonds stabilize the αB helix and shifts the equilibrium towards oligomerization for the L252P mutant. A previous study^42^ reporting molecular dynamics (MD) simulations on the L252P MyD88 TIR domain suggested that the L252P mutation promotes homo-oligomerization by reducing αC helix flexibility by a salt bridge formed by residues D225 and K258. However, we did not observe this interaction in either the WT or L252P structures. Instead, we propose that the L252P mutation restrains the conformational flexibility of the TIR domain by stabilizing the αB helix via K250, which might potentially also impact the BB loop conformation. Interestingly, the CD loop has been reported to undergo post-translational modification, specifically phosphorylation at the activating residue S244.^43,44^ This may highlight the important role of the CD loop in allosterically controlling MyD88^TIR^ homo-oligomerization.

Overlay of the WT MyD88^TIR^ ring structure and the L252P mutant spiral structure uncovers subtle rearrangement of the subunit interfaces due to the different CD loop conformations, which drives the transition from a closed ring to a spiral-like structure (Figure 6G). When one subunit of the WT ring (P1) is structurally aligned with that of the L252P spiral, the next subunit (P2) within the same filament and P1’ and P2’ subunits in the opposing filament all twist slightly at the four-subunit junction as if a “gear” there has been rotated (Figure 6H). Thus, by modulating both intrastrand and interstrand interactions, the small shift in L252P, repeated across protomers in the filament, leads to the formation of a spring-like structure instead of the closed ring seen in the WT (Video S4).

We wondered if autoactivation of L252P is associated with DNA binding. To address this question, we imaged MyD88 KO iBMDMs reconstituted with L252P. Strikingly but consistent with autoactivation, L252P iBMDMs transfected with buffer already displayed spontaneous MyD88 puncta, which colocalized with cytosolic dsDNA (Figure 6I). Transfection with DNA further enhanced MyD88 and dsDNA puncta in L252P iBMDMs (Figure 6I). These data support, although not demonstrate, a role of endogenous cytosolic DNA (e.g. from DNA damage or even cell division) in L252P activation.

## Discussion

Our findings uncover a previously unrecognized role of MyD88, revealing that this canonical adaptor may engage directly with cytosolic dsDNA that activates innate immunity independent of TLR engagement or signaling. Our high-resolution cryo-EM structures of the WT MyD88^TIR^ ring and the L252P spiral both reveal bound dsDNA. A previous study^45^ has reported that a plant TIR binds dsDNA to execute its function as a cAMP/cGMP synthetase. We demonstrated direct dsDNA binding of MyD88^TIR^ *in vitro*. Similarly, the mammalian TIR domain-containing protein SARM1 has been shown to directly bind dsDNA via its TIR domain, triggering its NAD^+^-degradation activity to promote cell death and chemotherapy-induced neuropathy.^46^ The DNA-binding defective MyD88 mutant K2E is incapable of DNA transfection-induced inflammatory cytokine expression while still capable of mediating TLR activation. Similarly, residues mediating the assembly of the double-layered TIR ring are required for dsDNA-induced pro-inflammatory cytokine expression, establishing the importance of the observed structure (Figure 7).

**Figure 7.**
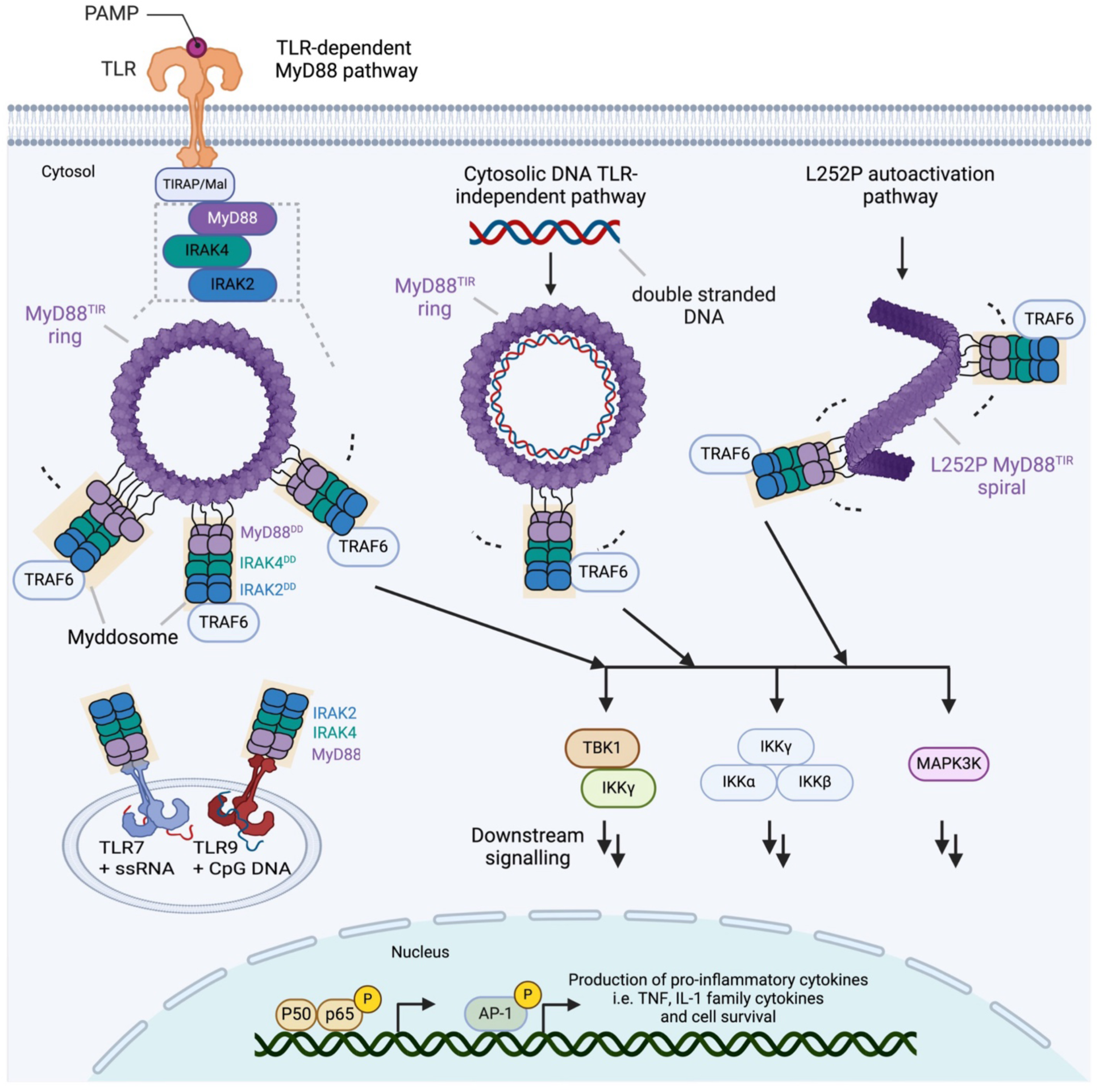
Model of MyD88-dependent NF-κB activation under different conditions. In the TLR-dependent pathway, MyD88 activation begins with dimerization of TLRs on the cell surface or in endosomes upon binding pathogen-associated molecular patterns (PAMPs). This interaction recruits MyD88 to the plasma membrane through TIRAP, enabling MyD88 molecules to assemble into Myddosomes initiated by the oligomerization of the MyD88^TIR^ domains. During this assembly process, the DD interactions promote recruitment of downstream kinases IRAK4, IRAK1 and IRAK2 via interactions with the MyD88 DD. This Myddosome formation activates the kinases and initiates a signaling cascade leading to NF-κB pathway activation, which ultimately triggers the expression of pro-inflammatory cytokines. In the dsDNA-dependent pathway, cytosolic detection of foreign or self-derived dsDNA triggers MyD88^TIR^ to organize into double-layered ring structures, with dsDNA bound to the concave ring surface. In cells with somatic mutations expressing the L252P MyD88, MyD88 shows autoactivation through spontaneous spiral formation, potentially leading to constitutive NF-κB activation. The schematic was created using BioRender.

Our unbiased RNA-seq experiments provided strong genetic evidence that MyD88 is a major contributor to transcriptional regulation induced by cytosolic DNA, controlling upregulation of genes in many pathways related to immune activation and host defense. Quite surprisingly, the number of DEGs cooperatively regulated by MyD88 is comparable to that observed with cGAS, the established cytosolic DNA sensor.^32,33^ Among all genes upregulated by dsDNA, most of them are co-regulated by MyD88 and cGAS, while the remaining are regulated by either MyD88 or cGAS. Generally, cGAS and MyD88 are most prominent for their roles in interferon pathway genes and inflammatory pathway genes, respectively. Given the co-dependence of most cytosolic DNA-induced genes on MyD88 and cGAS, it is tempting to speculate that stimulation of antitumor immunity requires activation of both cGAS-STING and MyD88 pathways. Conversely, suppressing autoimmunity might need to inhibit both cGAS-STING and MyD88 pathways.

MyD88 is ubiquitously expressed in different cell types and is evolutionarily ancient with conservation across a wide range of metazoans. It has been noted that cytosolic dsDNA can also activate NF-*κ*B, one of the most ancient and conserved immune pathways in animals, as well as the MAP kinase pathway, in normal and cancer cells but with unclear mechanisms.^33^ While many more details need to be resolved, our data are consistent with an intertwined relationship between cGAS and MyD88.

Our experimental data implicate a mechanistic model for how cytosolic dsDNA present during bacterial or viral infections or under sterile cellular stress activates the MyD88 signaling pathway independent of TLR signals using double-layered ring structures (Figure 7). We speculate that the kind of MyD88 assembly is also induced by TLR or IL-1R activation, which is supported by a recent study reporting analogous WTMyD88^TIR^ ring and cylindrical filament structures formed *in vitro* without dsDNA.^47^. In addition, we uncovered that Myddosome puncta formed by MyD88 and dsDNA are also barrel shaped, like the micron-sized puncta of Myddosome involved in TLR signaling.^31^ We surmise that the much smaller MyD88 rings or spirals could be the building blocks for these larger barrels in TLR, IL-1R, and DNA-induced MyD88 pathways, likely by condensate formation or other mechanisms.

Our study also provided a structural rationale of how the L252P mutation, frequently observed in a subset of lymphoma patients, triggers autoactivation of the MyD88 signaling pathway (Figure 7). Our data suggest that this autoactivation is likely dependent on dsDNA. Because lymphoma cells or other cancer cells may have more DNA damage or exhibit other features for cytosolic exposure to dsDNA, spontaneous NF-κB activation may occur in these cells, leading to survival signaling and cancer promotion. Collectively, our work highlights an unexpected role of MyD88 in dsDNA signaling and implicates the molecular architecture for MyD88-mediated pathway in TLR/IL-1R signaling, dsDNA sensing, and cancer.

## Methods

### Construct expression and purification

For structural studies, the human MyD88 TIR domain (155-296) was cloned into a pDB.His.MBP vector. His-MyD88 TIR domain and full length His-MyD88 were cloned into a pet28a vector. All mutations were introduced using a Q5 polymerase mutagenesis protocol. The construct was transformed into BL21-CodonPlus(DE3)-RIPL cells and plated on LB-Agar supplemented with kanamycin (30 µg/mL). A single colony was picked and grown overnight in Luria broth (LB) media supplemented with kanamycin as a pre-culture. For expression, LB media supplemented with kanamycin was inoculated with 1% pre-culture and grown to an OD_600_ of 0.6–0.8. The culture was then incubated for 30 minutes at 4 °C before adding 1 mM isopropyl β-D-1-thiogalactopyranoside (IPTG) to induce expression overnight at 18 °C. Bacterial cells were harvested by centrifugation at 3,000 g for 20 minutes at 4 °C. The pellet was resuspended in Buffer A (40 mM HEPES pH 7.0, 150 mM NaCl, 5 mM imidazole). Lysis was performed by sonication (1 second on, 5 seconds off, 3 minutes total on, 42% power, Branson) and subsequently centrifuged at 40,000 g for 45 minutes at 4°C. The soluble fraction was incubated with Ni-NTA resin and purified according to the manufacturer’s recommendation. Elution fractions containing MyD88 protein were concentrated using an Amicon Ultra 50 kDa MW cutoff and loaded on a Superose 6 Increase (Cytiva) size exclusion chromatography (SEC) column in SEC buffer (40 mM HEPES pH 7.0, 150 mM NaCl). Protein quality was assessed after each purification step by SDS-PAGE.

### Negative stain grid preparation and analysis

A 10 µL sample of the SEC void fraction containing MyD88 was loaded onto a glow-discharged negative stain grid (Formvar/Carbon Film 10/1 nm on Square 400 mesh) for 60 seconds, and excess protein was blotted onto filter paper. Grids were washed twice with Buffer A and stained with a 1% phosphotungstic Acid (PTA) solution for 30 seconds, blotted, and air-dried for a few minutes. The grids were imaged on a JEOL JEM1400 Transmission Electron Microscope operating at 120 keV.

### Cryo-EM grid preparation and data collection

The purified MyD88^TIR^ WT rings and L252P spirals from the void fractions were concentrated to 1 mg/mL and 2 mg/mL, respectively. The concentrated samples (4 µL) were loaded onto a glow-discharged Quantifoil (R 1.2/1.3 Au 400-mesh, Electron Microscopy Sciences), blotted for 3–5 seconds under 100% humidity at 4 °C, and plunged into liquid ethane with a Mark IV Vitrobot (ThermoFisher). All grids were screened before collection for ice thickness, particle quality, and distribution. WT samples were collected at the National Cancer Institute Cryo-EM facility, with a Titan Krios microscope (ThermoFisher) at an acceleration voltage of 300 keV equipped with a BioQuantum K3 Imaging Filter (slit width 20 eV), and a K3 direct electron detector (Gatan) operating in counting mode at 105,000× magnification (0.835Å pixel size). The L252P samples were collected at the S2C2 Stanford-SLAC Cryo-EM facility with a Titan Krios microscope (ThermoFisher) at an acceleration voltage of 300 keV equipped with a SelectrisX energy filter (slit width 20 eV), and a Falcon 4i direct electron detector (ThermoFisher) operating in counting mode at 105,000× magnification (0.95Å pixel size).

### Cryo-EM data processing

All data were processed using support for the infrastructure from SBGrid Consortium.^48^ Both data sets for WT MyD88^TIR^ and L252P MyD88^TIR^ were processed similarly. The raw movies were imported into CryoSPARC^49^ version 4.5.3. Beam-induced motion was corrected using Patch Motion, and contrast transfer function (CTF) parameters were estimated with PatchCTF. Images with an estimated CTF resolution worse than 6 Å were discarded, leaving 7,946 good images for the WT MyD88^TIR^ and 32,194 good images for the L252P MyD88^TIR^.

For WT MyD88^TIR^, particle coordinates were identified using a custom-trained Topaz^50^ model, resulting in the extraction of 3,792,682 particles with a box size of 400x400 and Fourier-cropped to 100x100 pixels. After several rounds of 2D classification, 391,591 particles with features characteristic of the MyD88^TIR^ ring segment were selected for further processing. An initial model was generated from the selected particles, and two junk models were created from discarded particles. These models served as references for multiple rounds of heterogeneous refinement (k=3, number of final full iterations=20). After each round, particles in the good class were selected for further rounds of heterogenous refinement, continuing until no more junk classes remained.

A total of 132,657 particles were re-extracted with a 400x400 box size and sequentially refined using homogeneous and heterogeneous refinement with C2 symmetry applied (window dataset (real space) *false*, minimize over per particle scale *true*), resulting in a 3.7 Å reconstruction. Following global and local CTF refinement, duplicate particle removal (separation distance < 20 Å), and 2D classification, 90,711 particles were further refined using non-uniform refinement to yield a 3.7 Å map with improved quality. Local refinement with a mask focused on the central octamer (2 layers of 4 subunits each) and C2 symmetry improved the resolution to 3.6 Å. To enhance the DNA density, 3D classification (k=30) with a DNA-focused mask produced a lower-resolution map with better DNA density. The final map was automatically sharpened in CryoSPARC using a soft mask and a B-factor of -105.8 Å^2^.

For the L252P MyD88^TIR^ dataset, 6,949,592 particles were initially extracted with a box size of 400x400 and Fourier-cropped to 100x100 pixels. After several rounds of 2D classification, 230,088 particles displaying features of the MyD88^TIR^ ring segment were selected for further processing through heterogeneous refinement and non-uniform refinement. Finally, 78,342 particles were refined locally with a soft mask around the central octamer and C2 symmetry applied, resulting in a 3.4 Å reconstruction. The final map was automatically sharpened in CryoSPARC using a soft mask and a B-factor of -100.5 Å^2^.

To process free DNA observed on the grid, we utilized the WT dataset. Initially, particle coordinates were determined using filament tracer. A total of 7,636,665 particles were extracted with a box size of 300x300 pixels, Fourier-cropped to 100x100 pixels, and subjected to multiple rounds of 2D classification (k=100). From these, 1,674,497 high-quality particles were selected for initial model generation. As described above, two junk models were generated from the discarded particles, serving as references for subsequent rounds of heterogeneous refinement. Subsequently, 109,455 particles were re-extracted with a box size of 300x300 pixels and refined using helical refinement, without applying any symmetry constraints. This iterative process resulted in a reconstruction with a resolution of approximately 6 Å. The final map was automatically sharpened in CryoSPARC using a soft mask and a B-factor of -100 Å^2^.

### Model building and figure preparation

To model the WT and L252P MyD88^TIR^ structures, the MicroED structure of MyD88^TIR^ (PDB ID: 7BEQ)^23^ was fitted as rigid body into the respective map using ChimeraX.^51^ A starting model for the dsDNA was generated by inputting a 60 bp herpes simplex virus 1 (HSV-60bp) reverse and forward strand DNA sequence into Alphafold3^52^, and fitted into the MyD88^TIR^ maps using ChimeraX.^51^ The models were iteratively refined in iSOLDE^53^ and Phenix^54^ until convergence was reached. Model validity was assessed in Phenix using MolProbity^55^ and EM Ringer score.^56^ ChimeraX was used to generate structural representations.

### Electrophoretic mobility shift assay (EMSA)

DNA oligos were purchased from Integrated DNA Technologies (IDT). Forward primers were modified by the addition of a Cy3 fluorescent probe. Forward and reverse primers were resuspended in 10 mM HEPES pH 7.0 and 50 mM NaCl, mixed to a 1:1 molar ratio and heated at 95 °C for 3 minutes in a heat block. The heat block was then left to cool down to room temperature allowing the formation of dsDNA.

Recombinant MBP-MyD88^TIR^ monomers at the indicated concentrations were incubated at 37 °C for 2 hours with 0.5 nM Cy3 labelled dsDNA in 40 mM HEPES pH 7.0, 150 mM NaCl, 0.2% BSA, and 5% glycerol. Bis-Tris 3-12% polyacrylamide electrophoresis gel (NativePAGE™) were pre-ran for 45 minutes at 4 °C in 0.5x running buffer, and the reactions were then loaded and ran for 75 minutes. Gels were quickly washed in running buffer and imaged by Cy3 fluorescence using a Typhoon™ laser scanner.

### Fluorescence size-exclusion chromatography (FSEC)

100nM of recombinant His-MyD88^TIR^ monomers were incubated at 37 °C for 1 hours with 20 nM Cy3 labelled dsDNA in 40 mM HEPES pH 7.0, 150 mM NaCl, and 5% glycerol. Sample were analysed by FSEC.

### Cell culture

Wild-type, cGAS knockout and MyD88 knockout immortalized bone marrow-derived macrophages and L929 cells were kindly provided by Dr. Jonathan Kagan. Endogenously GFP-tagged MyD88 expressing iBMDM cells were previously described^31^ and also kindly provided by Dr. Jonathan Kagan. All cells were cultured in DMEM containing 10% heat-inactivated and sterile-filtered fetal bovine serum (FBS), penicillin and streptomycin, and supplements of 2 mM L-glutamine and 1 mM sodium pyruvate. All cell lines were grown under sterile and standard cell culture conditions (humidified atmosphere, 5% CO_2_) and routinely tested for mycoplasma. All cells were passaged 1:10 every 2 days to prevent overcrowding and media acidification.

### Mice and bone-marrow-derived macrophage (BMDM) differentiation

MyD88^-/-^ knockout mice model was previously described.^57^ C57BL/6J mice were used as a control. All mice were maintained in a pathogen-free animal facility at the Boston Children’s Hospital. For obtaining the bone marrow-derived macrophages, sex and age matched 6- to 12-week-old mice were selected for wild-type and MyD88^-/-^ mice. Primary bone marrow-derived macrophages from mice were cultivated 7 days in DMEM supplemented with 10%FBS, 30% L929-conditioned media, 1% penicillin and streptomycin and 1% non-essential amino acids. The differentiation medium was replenished every 2-3 days and differentiation was verified by double F4/80 and CD11b double staining by flow cytometry.

### Intracellular nucleic acid delivery

Wild-type, cGAS or MyD88 knockout iBMDMs were seeded into 6-well plates at 1.5x10^6^ cells/well density 24 hours prior to the delivery of the respective nucleic acids. Indicated lengths of dsDNA or poly I:C (high molecular weight, cat.no. tlrl-pic-5) were prepared at 0.1 mg/mL concentration. Double-stranded DNA of 2000 and 7000 bp was amplified by PCR from a custom endotoxin-free pCDNA 3.1 vector. Cells were transfected using Lipofectamine 3000 (Invitrogen, cat.no. L3000001) according to manufacturer’s instructions. Briefly, 2.5 µg of respective nucleic acid was diluted in 125 µl of OPTIMEM. For each reaction, 5 µl of P3000 reagent was then added to the mixture. In a separate tube, 7.5 µl of Lipofectamine 3000 was diluted in 125 µl of OPTIMEM. After 5 minutes incubation at room temperature, nucleic acid-containing mixture was added to the lipofectamine and incubated for 15 minutes at room temperature. Nucleic acid-lipofectamine complexes were then added to the cell media in a drop-wise manner.

### HSV-1 infection

On the final day of differentiation, primary BMDMs were seeded into 12 well plates at 1x10^6^ cells/well density. Cells were infected with HSV-1 KOS strain (HSV_KOS)^58^ at 100 MOI or with HSV-1 mutant 7134 strain (HSV_ΔICP0)^59^ at 10 MOI in DMEM plain media for 18 hours,. Supernatants were collected for ELISA and the remaining cells harvested for subsequent RNA extraction.

### Cell stimulations

Cells were harvested 6 hours after nucleic acid delivery. The two-step activation of NLRP3 inflammasome was performed as previously described.^60^ Briefly, iBMDMs were seeded to 6-well plate at 1.5 x 10^6^/well density 24 hour prior to the treatment. Cells were primed with 1 µg/mL of LPS (Invivogen, cat.no. tlrl-pb5lps) for 4 hours and stimulated with 20 µM of nigericin (Sigma-Aldrich, cat.no. N7143) for 30 minutes. For TLR activation, cells were stimulated with the following TLR1/2 (1 μg/ml Pam3CSK4), TLR7/8 (1 μg/ml, R848), TLR4 (0.1 μg/ml, LPS-B5 for 18 hours, or TLR9 (2 μM ODN1826) agonists.

### Lentiviral production and transduction to iBMDMs

293T cells were seeded into 10 cm plate in DMEM media containing 10% FBS without antibiotics at 70% density 20-24 hours prior the transfection. 8 µg of the indicated transfer vector pLV was diluted in 1 ml OPTIMEM together with 4 µg of packaging vector psPAX2 and envelope plasmid pMD2.G. 80 µl of Lipofectamine 2000 was then added to the mixture and incubated for 20 minutes at room temperature. The transfection mixture was the add to the 293T cell media in a drop-wise manner. Cell media was change at 6 hours post transfection and lentiviral supernatants were harvested 72 hours later. Supernatants were then sterile filtered using 0.45 µm PES filter and centrifuged at 1500 RPM for 5 minutes at room temperature to remove any cell debris. Supernatants were then mixed with a lentiviral concentrator (Lenti-X-Concentrator, Takara), at 1:3 ratio and incubated at 4 °C overnight. Lentiviruses were then concentrated using centrifugation at 1500xg for 45 minutes at 4 °C and lentivirus containing pellets were resuspended in 1 ml DMEM plain media.

Lentiviruses were immediately transduced to iBMDM cells using spinfection. Briefly, 0.75 x10^6 cells were seeded to each well of 12-well plate and 120 µl of concentrated virus was then added. Polybrene was then added to the mixture at 5 µg/ml concentration. Cells were spinfected at 932xg for 60 minutes at 37 °C and then placed in the incubator. 48 hours post transduction, 7.5 µg/ml puromycin was added to the cells to select the transduced population.

### CRISPR/Cas9-mediated knockout

CRISPR knockout was performed utilizing pLentiCRISPRv2 plasmid.^61^ The gRNA containing plasmids were transfected into 70-80% confluent 293T cells with the following reaction mixture: 8 µg of the indicated gRNA-containing CRISPR vector was diluted in 1 ml OPTIMEM together with 4 µg of packaging vector psPAX2 and envelope plasmid pVSVG. 80 µl of Lipofectamine 2000 was then added to the mixture and incubated for 20 minutes at room temperature. The transfection mixture was the add to the 293T cell media in a drop-wise manner. Cell media was change at 6 hours post transfection and lentiviral supernatants were harvested 48 hours later. Supernatants were then sterile filtered using 0.45 µm PES filter and centrifuged at 1500 RPM for 5 minutes at room temperature to remove any cell debris. Supernatants were then mixed with a lentiviral concentrator (Lenti-X-Concentrator, Takara), at 1:3 ratio and incubated at 4 °C overnight. Lentiviruses were then concentrated using centrifugation at 1,500xg for 45 minutes at 4 °C and lentivirus containing pellets were resuspended in 1 ml DMEM plain media. Lentiviruses were immediately transduced to iBMDM cells using spinfection. Briefly, 0.75 x10^6 cells were seeded to each well of 12-well plate and 120 µl of concentrated virus was added. Polybrene was then added to the mixture at 5 µg/ml concentration. Cells were spinfected at 932xg for 60 minutes at 37 °C and then placed in the incubator.

48 hours post-transduction cells were selected with 7.5 µg/ml puromycin for 72 hours. Single cell clones were obtained using single-cell sorting. Genomic DNA was extracted from the cells and genomic area 400-500 bp spanning around the gRNA-target site was amplified via PCR, gel purified and sequenced. The sequencing reads were compared with the wild-type sequence using ICE CRISPR Analysis tool (Synthego) to determine the knockout containing clones. Subsequently, the knockout was verified by immunoblotting.

### Indirect immunofluorescent (IF) imaging

IF imaging was performed as previously described.^61^ Briefly, cells were seeded into coverslips containing 6-well plates and stimulated as indicated. 2-3 hours post dsDNA transfection, cell media was aspirated and cells were washed twice with PBS. Cells were then fixed in 4%PFA for 15 minutes at room temperature. PFA was then washed away 3-times with PBS. Cells were permeabilized with 0.1% Triton_X_100 for 15 minutes at room temperature. Permeabilization buffer was then washed away 3 times with PBS washes and cells were blocked with 6%BSA in PBS for 1 hours at room temperature. Coverslips were embedded in dsDNA marker primary antibody at 1:100 dilution in 6%BSA in PBS solution at 4 °C for overnight. Coverslips were washed with PBS for 3 times and were embedded in anti-mouse Alexa Fluor 647 nm for 1 hour at room temperature in 6%BSA in PBS. Coverslips were then washed 3 times with PBS and samples were incubated 20 µM of Hoechst staining solution for 15 minutes at room temperature. Coverslips were then embedded in approximately 10 µl of Vectashield.

Immunofluorescence images were acquired using a spinning disk confocal microscope with a 63× oil immersion objective. Z-stacks (13–15 slices per field) were collected to capture the full depth of the cells. Maximum intensity projections were generated using ImageJ to visualize and analyse the data. To verify signal specificity and exclude the signal bleed through between channels, the IF staining procedure was in parallel performed under no primary or secondary antibody and no Hoechst conditions.

### Western blotting

Cell media was aspirated, and cells were washed twice with ice-cold PBS. 150 µl RIPA buffer containing EDTA-free Protease Inhibitor (Sigma-Aldrich, cat. no. 11836170001) and PhosSTOP (Sigma-Aldrich, cat. no. 4906837001) cocktails were added to each well. Cells were then scraped with the lysis buffer and transferred into 1.5 mL tubes. Lysates were incubated at 4 °C for 20 minutes with end-to-end rotation and spun down at the highest setting in an Eppendorf centrifuge for 10 minutes. Supernatants were obtained as whole cell lysates.

The whole cell extracts were mixed with Laemmli Buffer and separated by electrophoresis using 4-20% gradient gels. Proteins were transferred and immobilized onto a nitrocellulose membrane using standard Towbin Buffer containing 20% methanol for 2 hours at constant 220 mA. For blocking, membranes were treated with 3% BSA in TBST for anti-IL-1β, and with 5% non-fat milk in TBST for all other antibodies. Membranes were incubated with the respective primary antibodies overnight at 4 °C, washed three times (5 minutes each) with TBST, and incubated with the corresponding isotype specific HRP-conjugated secondary antibody in 1% non-fat dry milk containing TBST for 2 hours at room temperature. Membranes were then washed three times (5 minutes each) with TBST, and chemiluminescent signals were visualized using standard ECL solution.

### Antibodies

The following primary antibodies were used for immunoblotting: Anti-IKKα + IKKβ phospho S176+S177 (Abcam, cat.no. ab194528, 1:1000 dilution in 1% non-fat dry milk in TBST), Anti-phospho-p65 (Ser536) (Cell Signaling Technology, cat. no. 3031, 1:2000 dilution in 1% non-fat dry milk in TBST), Anti-phospho STING (Ser365) (Cell Signaling Technology, cat. no. D8F4W, 1:1000 dilution in 3% BSA in PBST), Anti-IL-1β (R&D Systems, cat. no. AF-401-SP, 1:1000 dilution in 5% BSA in TBST), Anti-GAPDH (Proteintech, cat. no. 60004-1-IG, 1:10000 dilution in 1% non-fat dry milk in TBST), Anti-beta-actin (Sigma-Aldrich, cat. no. A3854, 1:10000 dilution in 1% non-fat dry milk in TBST), Anti-MyD88 (Cell Signaling Technology, cat. no. D80F5, 1:1000 dilution in 1% non-fat dry milk in TBST), Anti-p65 (Cell Signaling Technology, cat.no. L8F6, 1:1000 dilution in 1% non-fat dry milk in TBST), rabbit anti-p-p38α (Thr180/Tyr182) (CST, #4511, 1:1000 in 1% non-fat dry milk in TBST), rabbit anti-pERK1/2 (Thr202/Tyr204) (CST, #4370, 1:1000 in 1% non-fat dry milk in TBST). The following antibodies were used for immunofluorescence as primary antibodies: MyD88 (Biotechne, cat no. AF3109, at 1.7 μg/ml) and dsDNA marker antibody (Santa Cruz, sc-58749, at 1:100 dilution).

The following secondary antibodies were used for their corresponding isotype specific primary antibodies; Peroxidase AffiniPure^TM^ Goat Anti-Rabbit IgG (H+L) (Jackson ImmunoResearch, cat. no. 115-035-003), Mouse anti-goat IgG-HRP (Santa Cruz Biotechnology, cat.no. sc-2354), and Peroxidase AffiniPure^TM^ Goat Anti-Rabbit IgG (H+L) (Jackson ImmunoResearch, cat. no. 115-035-003), donkey anti-goat IgG (H+L) Cross-Adsorbed Secondary Antibody, Alexa Fluor^TM^ 647 (Thermo Fisher Scientific, cat. no. A-21447), donkey anti-mouse IgG (H+L) Cross-Adsorbed Secondary Antibody, Alexa Fluor^TM^ 568 (Thermo Fisher Scientific, cat. no. A10037).

### LDH release

Wild-type or MyD88 knockout iBMDMs were seeded into 6-well plates at 1.5x10^6^ cells/well density 24 hours prior to the treatment. For each cell type, quadruplicate wells were seeded for determination of the maximum LDH content. Cells were treated or transfected with the respective nucleic acids as indicated. The maximal LDH release wells were treated with 0.1% Triton-X-100 at final concentration for 10 minutes before collecting the samples. After treatment, cell supernatants were collected and spun at 1,500 RPM for 5 minutes to eliminate the floating cells. Supernatants were diluted to 1:20 in LDH storage buffer containing 200 mM Tris-HCl (pH 7.3), 10% Glycerol, 1% BSA. LDH levels were measured using LDH-Glo^TM^ Cytotoxicity Assay kit (Promega, cat. no. J2380) according to the manufacturer’s instruction. Percentage cytotoxicity was calculated according to the following formula: Percent Cytotoxicity=100 x (Experimental LDH-Medium Background) / (Maximal LDH Release Control-Medium Background).

### Gene expression analysis

Wild-type, MyD88 or cGAS knockout iBMDMs were treated as indicated. Total RNA was isolated using TRizol^TM^ Reagent (Invitrogen, cat. no. 15596026) according to the manufacturer’s instruction. In brief, cell supernatants were aspirated after indicated stimulations, and cells were washed twice with PBS. Cells were then directly harvested in 1 ml TRizol and incubated at room temperature for 2-3 minutes, followed by addition of 200 µl chloroform. After phase separation by centrifugation (12,000 g for 35-55 minutes at 4 °C), total RNA was precipitated from the aqueous layer with isopropanol. The RNA pellet was washed three times with 75% RNase-free ethanol, air-dried and resuspended in nuclease-free water.

For cDNA synthesis, a total of 1,000 ng RNA was used for each condition. High-Capacity cDNA Reverse Transcription Kit (Applied Biosystems, cat. no. 4368813) was used according to the manufacturer’s instruction to reverse transcribe the RNA into cDNA. The cDNA was subsequently diluted 1:20 in nuclease free water.

qRT-PCR was performed as described previously.^62^ Primers were designed according to MIQE guidelines (Minimum Information for Publication of qRT-PCR Experiments) using exon-intron spanning primers with 62 °C melting temperature. The DNA folding tool from the mfold server was used to exclude primers forming secondary structures. Primer efficiencies were determined using three serial dilutions of cDNA and only primers with efficiencies around 98-100% were chosen for further analysis. To determine fold induction of target genes, two reference genes with an M value lower than 0.5 were used, Hrpt1 and Gapdh respectively. For the detection of the qRT-PCR reaction, GoTaq qPCR Master Mix, 2X (Promega, cat. no. A600A) was used. For a single reaction, 5 µl of GoTaq qPCR Master Mix was mixed with 1 µl forward and reverse primer pairs (0.3 µM final concentration), 2 µl of nuclease free water and 2 µl of cDNA template. Samples were prepared in quadruplicates and reaction was run using CFX96 Touch Real-Time PCR Detection System (Bio-Rad) using the following setup: Stage 1: Activation at 50 °C for 2 minutes; Stage 2: pre-soak at 95 °C for 10 minutes; Stage 3: Denaturation at 95 °C for 15 seconds, Annealing at 60 °C for 1 minute; Stage-4: Melting Curve, 95°C for 15 seconds, 60 °C for 15 seconds, 95 °C for 15 seconds.

### RNA-seq

Cells were transfected with either dsDNA or annealing buffer control. Cells were harvested in TRizol^TM^ Reagent (Invitrogen, cat. no. 15596026) 6 hours post-transfection and the total RNA was extracted according to the manufacturer’s protocols. RNA quality and RNA Integrity Number (RIN) were determined using Agilent TapeStation High Sensitivity RNA. For total bulk mRNA sequencing, Next Generation Library Preparation was performed using KAPA mRNA Hyper Prep with 50 ng RNA for each condition according to the manufacturer’s instructions. Briefly, mRNA is pulled down using oligo-dt beads, and resulting mRNA is converted into cDNA. The resulting cDNA then became a library through adaptor ligation and post-PCR Cleanup. For the library sequencing, NextSeq2000 P3 (XLEAP-SBS, 100 cycles with 1.2 billion reads depth) kit was used with Illumina NovaSeq X Plus Sequencer instrument.

RNA-seq counts were generated by the nf-core rnaseq pipeline using Salmon.^63^ Downstream analyses were performed using R version 4.4.1 (2024-06-14). Counts were imported into R using DESeq2 version 1.46.9.^64^ Gene annotations were obtained from Ensembl. Plots were generated by ggplot2.^65^ Heatmaps were generated by pheatmap. Differential gene expression analysis of count data was performed using the Bioconductor R package, DESeq2. RNA-seq differential expression and gene-set enrichment analysis are available as Supplementary Tables 1 and 2, respectively.

### ELISA

Cells were stimulated with the indicated nucleic acids as described and cell supernatants were collected after the treatments. To remove the floating cells, supernatants were filtered, and cell debris were spun out at 1,500 RPM for 5 minutes. The levels of respective cytokines were determined following manufacturer’s instructions: mouse TNF DuoSet ELISA (R&D Systems, cat. no. DY410-05) and mouse IFN-β DuoSet ELISA (R&D Systems, cat. no. DY8234-05).

### Stimulated emission depletion (STED) microscopy

STED super-resolution microscopy of fixed samples was performed using an Abberior STEDYCON 2-colour STED microscope (Abberior Instruments). Imaging was performed using a Plan Apo Lambda ×100/1.45 oil-immersion objective (Nikon). The following settings were used: 16 bit bit-depth, 5× line averaging, ROI 500 × 500 px with a pixel size of 25 nm per px and a pinhole size of 64 µm. For imaging, 0.125 × 10^6^ cells were seeded on coverslips in 6-well plates 20-24 hours prior to the stimulation. Cells were then transfected with HSV-60bp for 6 hours to induce Myddosome formation and fixed with 4% methanol-free formaldehyde for 15 minutes at room temperature. Fixed cells were washed with PBS, permeabilized and blocked with PermQuench buffer (Thermo Fisher Scientific, cat no. 00-8333-56) for 30 minutes at room temperature. Fixed cells were co-stained with 1:100 diluted primary antibodies against the Flag-tag in MyD88–AGF and dsDNA with dsDNA marker for 1 hour at room temperature in PermQuench buffer. After three washes with PBS, cells were incubated with 1:100 diluted STAR Red- and STAR Orange-labelled secondary antibodies (Abberior) in PermQuench buffer for 1 h at room temperature. Coverslips were extensively washed with PBS, once with double-distilled H_2_O and mounted using ProLong Gold Antifade Mountant (P36930, Thermo Fisher Scientific) and left to cure overnight. The acquired data were processed by merging channels in Fiji (ImageJ) and adjusting contrast to enhance the visualization of Myddosomes, which were classified based on their appearance

## Data availability

The cryo-EM maps have been deposited in the Electron Microscopy Data Bank under the following accession codes: WT MyD88^TIR^ after non-uniform refinement (16mer, EMD-XXXXX), WT MyD88^TIR^ after the final local refinement (octamer, EMD-XXXXX), WT MyD88^TIR^ with enhanced density of the bound DNA after 3D classification (16 mer, EMD-XXXXX, provided as additional map), L252P MyD88^TIR^ after non-uniform refinement (16 mer, EMD-4 XXXXX), L252P MyD88^TIR^ after the final local refinement (octamer, EMD-XXXXX). Atomic coordinates of the final models have been uploaded to the Protein Data Bank with the accession codes: WT MyD88^TIR^ octamer (XXXX) and L252P MyD88^TIR^ octamer (XXXX).

## Acknowledgments

We thank Prof. Jonathan Kagan for wild-type, cGAS KO and MyD88 KO iBMDMs, L929 cells, and endogenously GFP- and FLAG-tagged MyD88 iBMDMs, and Prof. Greg Barton for WT and *Unc93b1* KO RAW 264.7 macrophages. We thank Arthur V. Hauenstein for his earlier work on this project. Cryo-EM data collection was conducted at the National Cancer Institute’s National Cryo-EM Facility at the Frederick National Laboratory for Cancer Research under contract 75N91019D00024, and the Stanford-SLAC Cryo-EM Center (S2C2) supported by the National Institute of General Medical Sciences (1R24GM154186). We thank the BPF Genomics Core Facility at Harvard Medical School (RRID:SCR_007175) for their expertise and instrument availability that supported this work. We thank Alex Bartlett, Emma Berdan and Shannan Ho Sui of the Harvard Chan Bioinformatics Core, Harvard T.H. Chan School of Public Health, Boston, MA, RRID:SCR_025373 for assistance with RNA-seq analysis. We thank the CITE (Core for Imaging Technology and Education) and MicRoN cores at Harvard Medical School for confocal and STED microscopy support, respectively. We thank Viraga Haridas (BCH FICR-PCMM Flow and Imaging Cytometry Facility) for assistance with single-cell sorting. This work was supported by National Institutes of Health grants R01AI050872 to HW and R21AI188827 to DMK, GM156406 to JAP and GM158136 to SPG. PF was supported by a Boston Children’s Hospital Office of Faculty Development/Basic & Clinical Translational Research Executive Committees Faculty Career Development Fellowship, ABT is supported by CRI-Irvington Postdoctoral Fellowship Program. L.S. is supported by the National Science Foundation Graduate Research Fellowship Program under Grant No. (DGE 2140743); any opinions, findings, and conclusions or recommendations expressed in this material are those of the author(s) and do not necessarily reflect the views of the National Science Foundation. L.S. is also supported by the Herchel Smith Graduate Fellowship.

## Author contribution

Conceptualization: PF, PG, ABT, HW. Methodology/Investigation: PF performed protein purification, biochemical assays, and cryo-EM data collection. PG and PF processed and analyzed cryo-EM data and helped with initial cell biology experiments. ABT performed cell biology experiments using HSV-1 infections, CRISPR KO, RNA-seq, gene expression analysis, and confocal microscopy. EG helped with gene expression and western blotting analyses. LS and SMV helped with RNA-seq data analysis. HSO helped with HSV-1 infections. ABT, JAP and SPG performed proteomics. PF, ABT, PG, HW and DMK wrote the manuscript with input from all authors.

---

**Figure S1.**
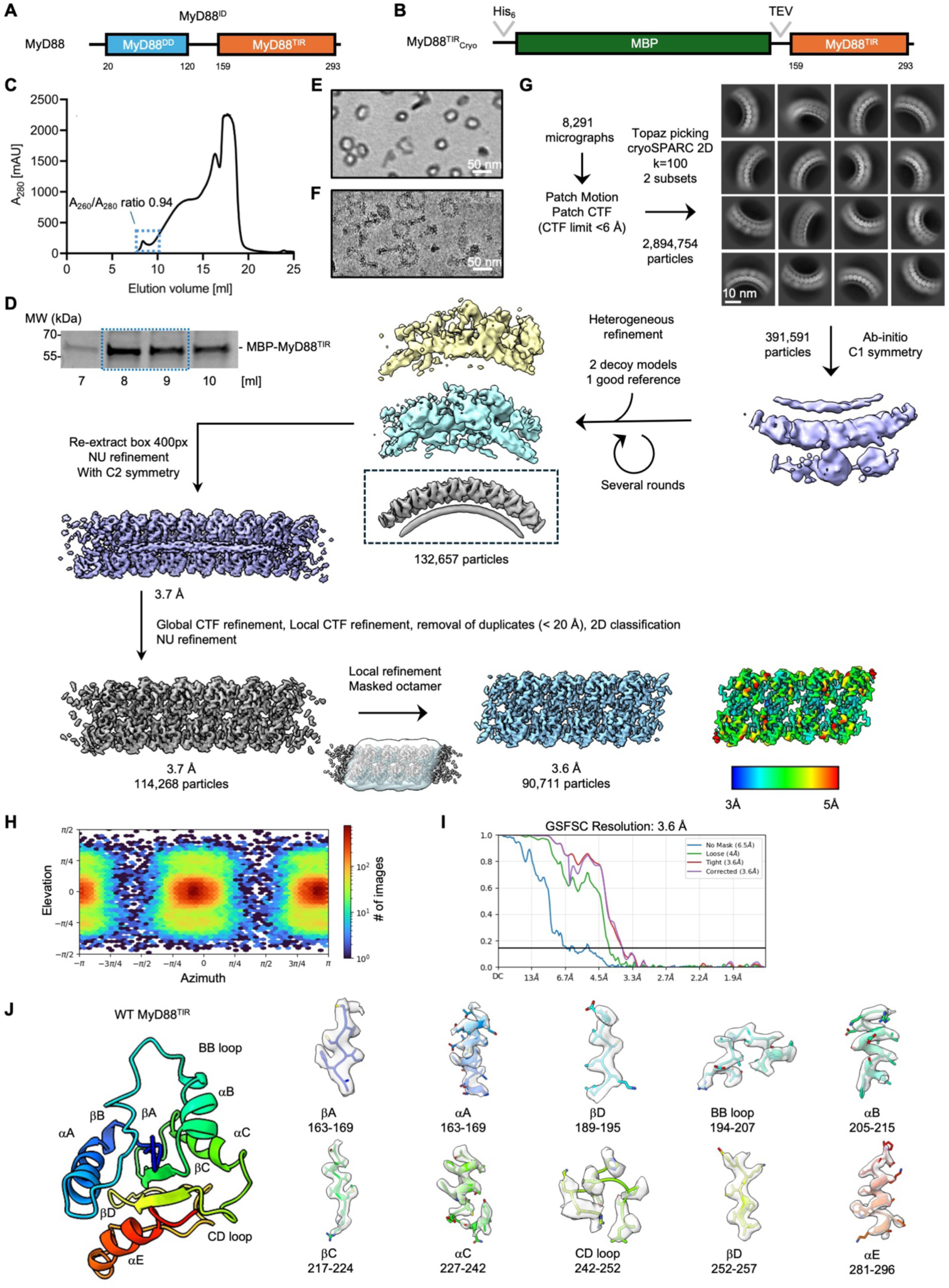
Cryo-EM processing workflow of the WT MyD88^TIR^ ring segment. **(A)** MyD88 domain organization. **(B)** Construct of MBP-tagged MyD88TIR used for structural determination. **(C, D)** Size-exclusion chromatography profile (C) and SDS-PAGE (D) from the final purification step of the MyD88TIR domain, with the indicated A260/A280 absorption ratio for the void peak. **(E)** Negative stain EM analysis of the MyD88^TIR^ domain sample from the void volume. Scale bar, 50 nm. **(F)** A cryo-EM image showing representative MyD88^TIR^ rings in vitreous ice. Scale bar, 50 nm. **(G)** Data processing workflow for the WT MyD88^TIR^ ring segment, detailing stages from initial micrograph collection to 2D classification and final local refinement. Scale bar for 2D class averages, 10 nm. **(H, I)** Orientation diagnostics from the final local refinement step, showing distribution of particle orientations (H) and the Fourier shell correlation (FSC) plot (I). **(J)** Map quality evaluation of the WT MyD88^TIR^ assembly. The cryo-EM map regions are displayed as transparent surface representations, with corresponding atomic models fitted into the density and shown in cartoon representation to highlight structural details.

**Figure S2.**
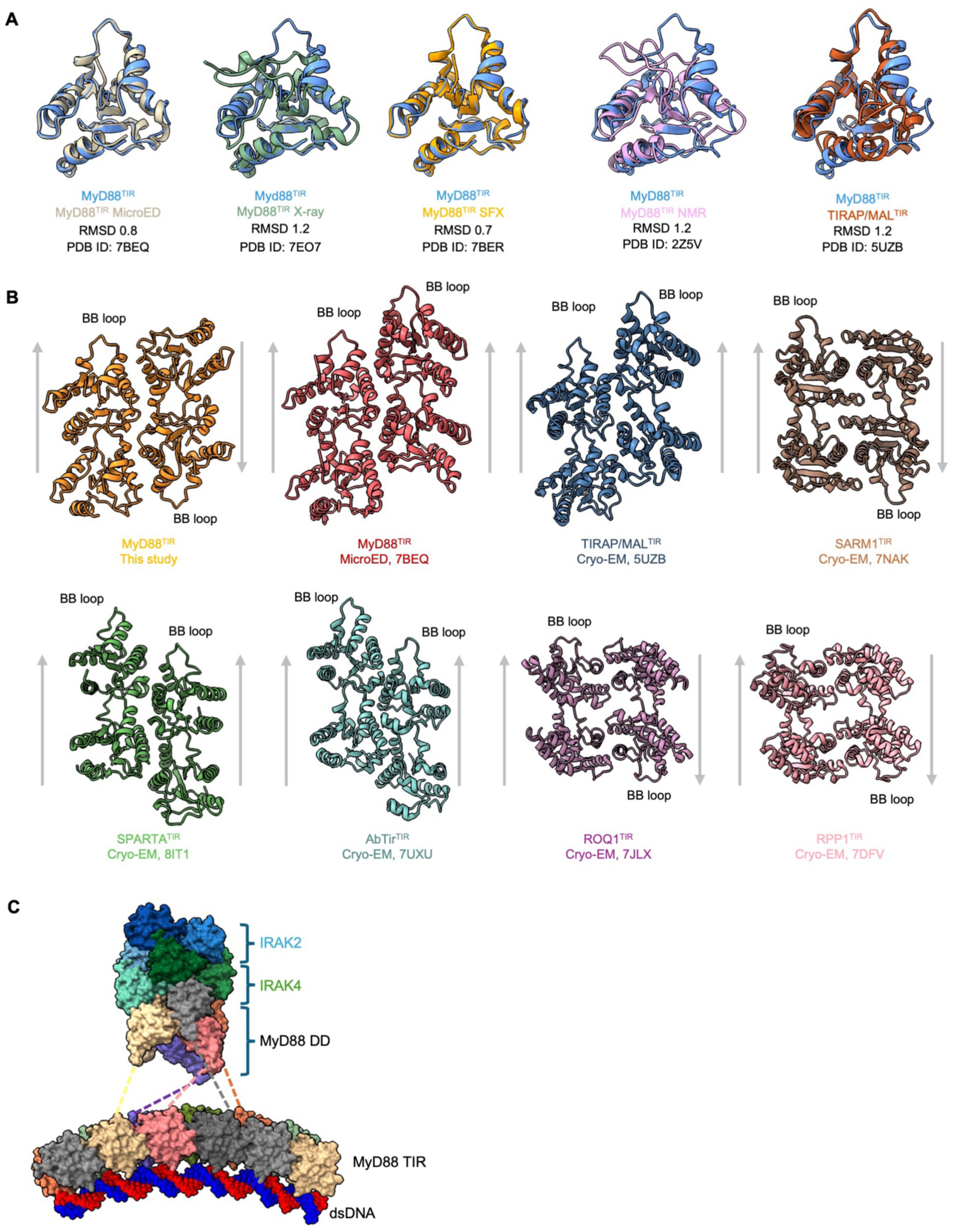
Structural overview and comparison of different oligomeric TIR structures. **(A)** Overlay of the MyD88^TIR^ protomer structure from the rings with other known structures of MyD88^TIR^ and TIRAP/Mal^TIR^. SFX: serial femtosecond crystallography. **(B)** Representation of different TIR filament or oligomeric structures in a cartoon representation of antiparallel MyD88 (orange, this study), parallel MyD88 (red, PDB ID 7BEQ), parallel TIRAP/MAL (blue, PDB ID 5UZB), parallel Ab^TIR^ (cyan, PDB ID 7UXU), parallel SPARTA (green, PDB ID 8IT1), antiparallel SARM1 (brown, PDB ID 7NAK), antiparallel ROQ1 (dark pink, PDB ID 7JLX) and RPP1 (light pink, PDB ID 7DFV). The grey arrows indicate the direction of the filament or structure from the EE surface to the BB loop. **(C)** Model of a MyD88^TIR^ ring segment with the helical oligomer of hexameric MyD88^DD^, tetrameric IRAK4^DD^, and tetrameric IRAK2^DD^ from the crystal structure of the MyD88:IRAK4:IRAK2 complex (PDB ID 3MOP).

**Figure S3.**
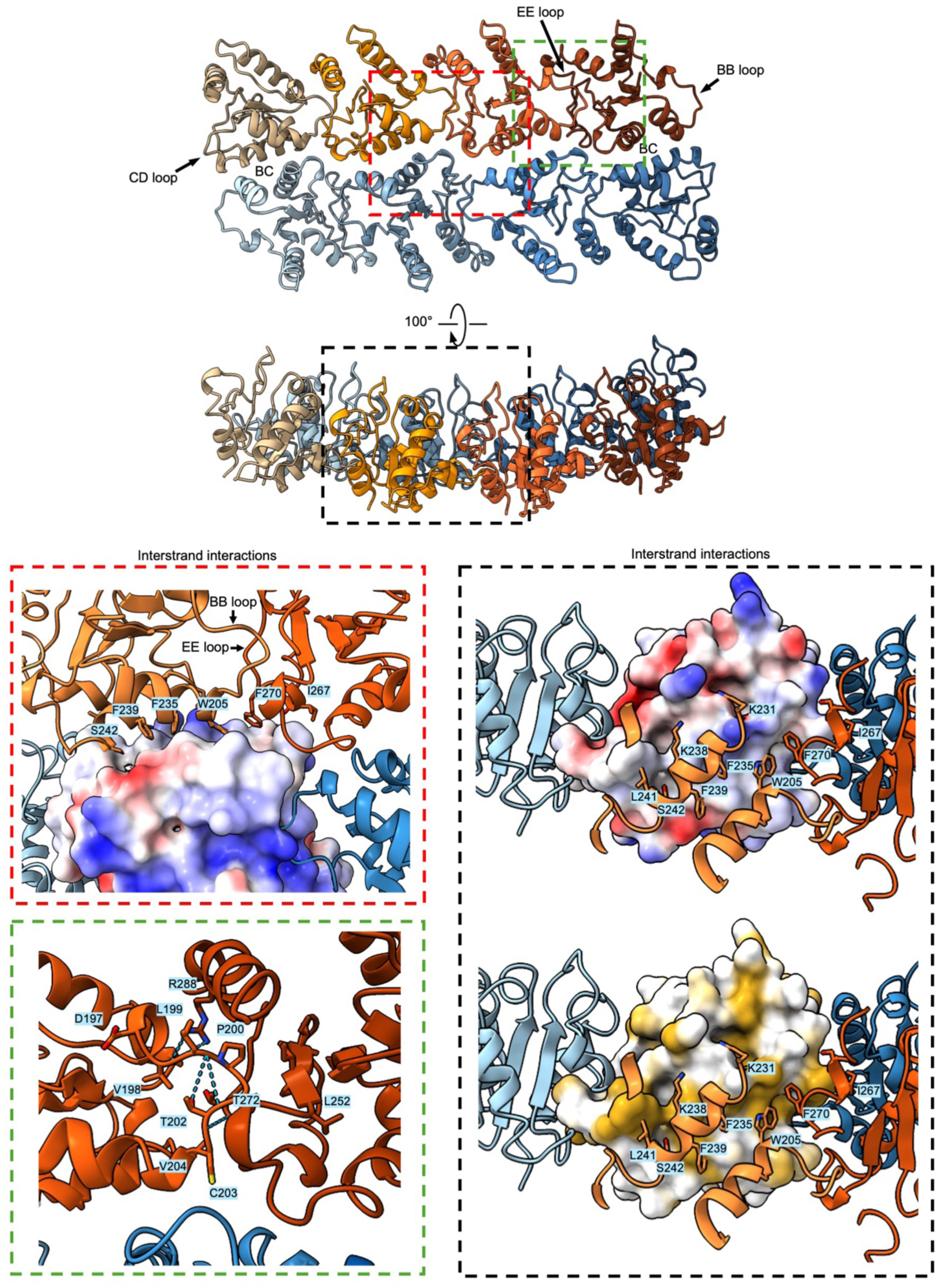
Interstrand interactions within the MyD88^TIR^ double-layered ring. Detailed interactions of the WT MyD88^TIR^ double-layered ring focusing on the unique interstrand interactions of the antiparallel TIR filaments. Close-ups show the interaction interfaces in different orientations. The surfaces are coloured either according to the coulombic surface potential (blue and red), or hydrophobicity (ochre) and hydrophilicity (white). Potential hydrogen bonds are indicated by dashed lines.

**Figure S4.**
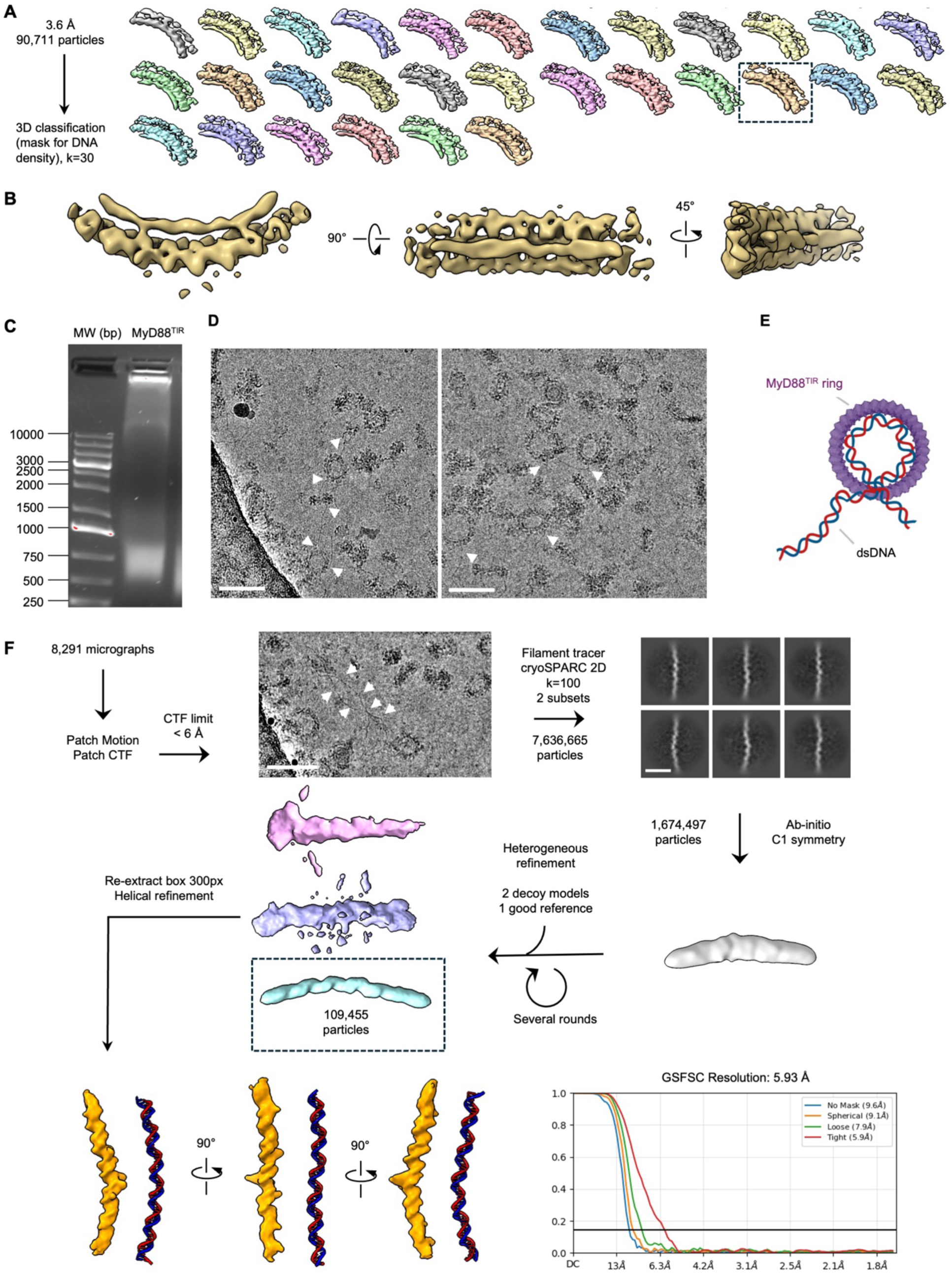
Processing workflow for the co-purified dsDNA. **(A)** Final particle set obtained from local refinement used as input for 3D classification with a soft mask applied to enhance density for the bound dsDNA region. **(B)** Views of the cryo-EM map from different orientations after 3D classification (A), highlighting improved density features of the bound dsDNA on the MyD88^TIR^ ring. **(C)** Purified MyD88^TIR^ sample used to prepare the cryo-EM grids was denatured and loaded onto a 1% agarose gel to visualize the size distribution of the co-purified dsDNA stained by SYBR® green. **(D)** Representative cryo-EM micrographs showing long dsDNA bound to MyD88^TIR^ rings. The white arrows indicate where the dsDNA enters or leaves the TIR rings. Scale bar: 100 nm. **(E)** Schematic representation showing how long dsDNA can bind to the MyD88^TIR^ ring. One TIR ring can bind approximately 240 bp of dsDNA. Excess dsDNA will protrude away from the TIR ring. The schematic was created using BioRender. **(F)** Data processing workflow for the free dsDNA observed on cryo-EM micrographs. Scale bars: micrographs, 100 nm; 2D class averages, 10 nm.

**Figure S5.**
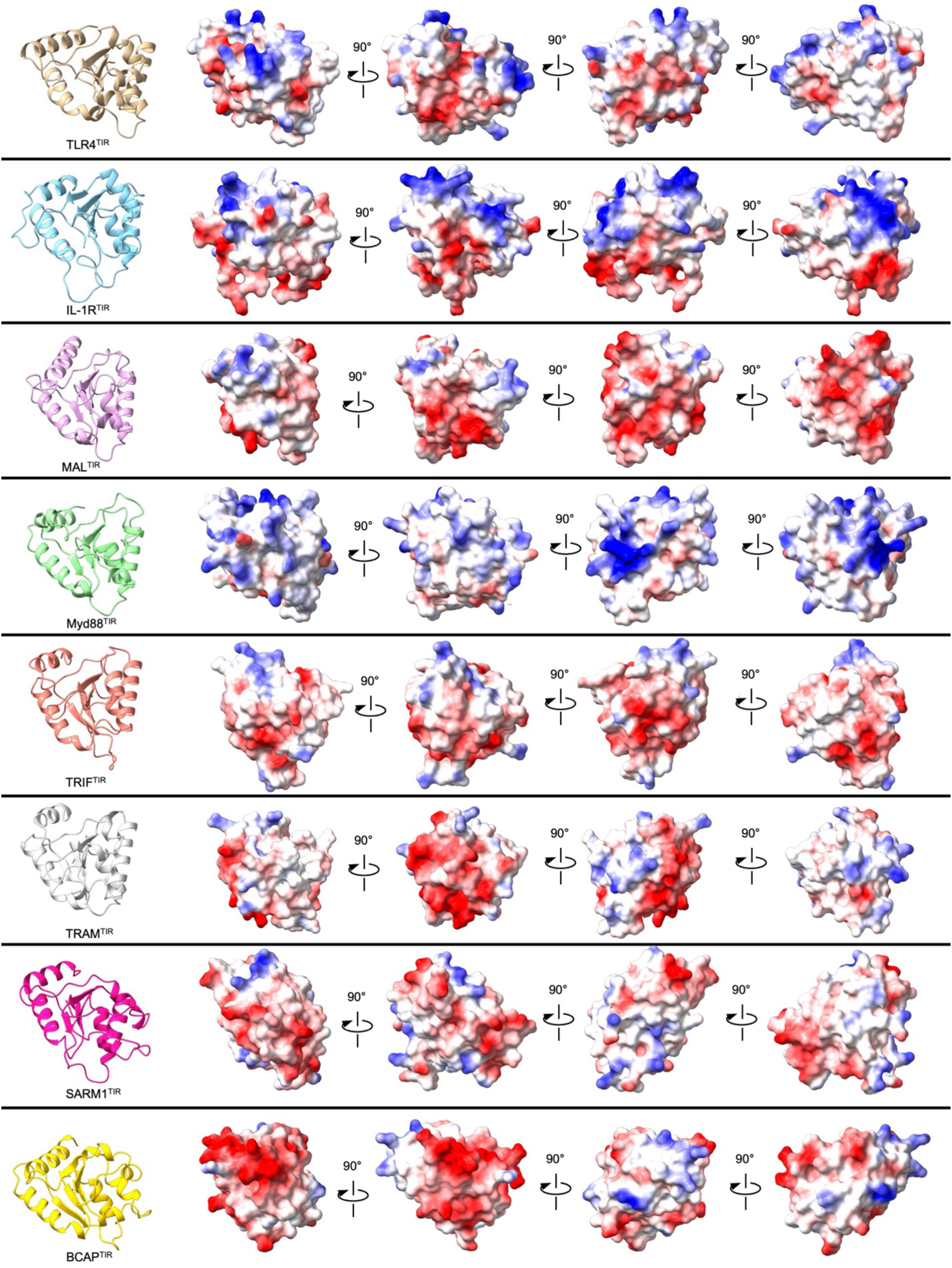
Electrostatic surface potential distribution of different human TIR domains. Each TIR domain is initially shown in a single orientation in cartoon representation. Additional views display the electrostatic surface charge distribution, with negatively charged regions highlighted in red and positively charged regions in blue. For TIR domains with available experimental structures, these structures were used in our analysis (PDB IDs: TIRAP/MAL, 5UZB; SARM1, 7NAK; BCAP, 5FOR). For TIR domains lacking experimental structures, we utilized predicted models generated by AlphaFold2.

**Figure S6.**
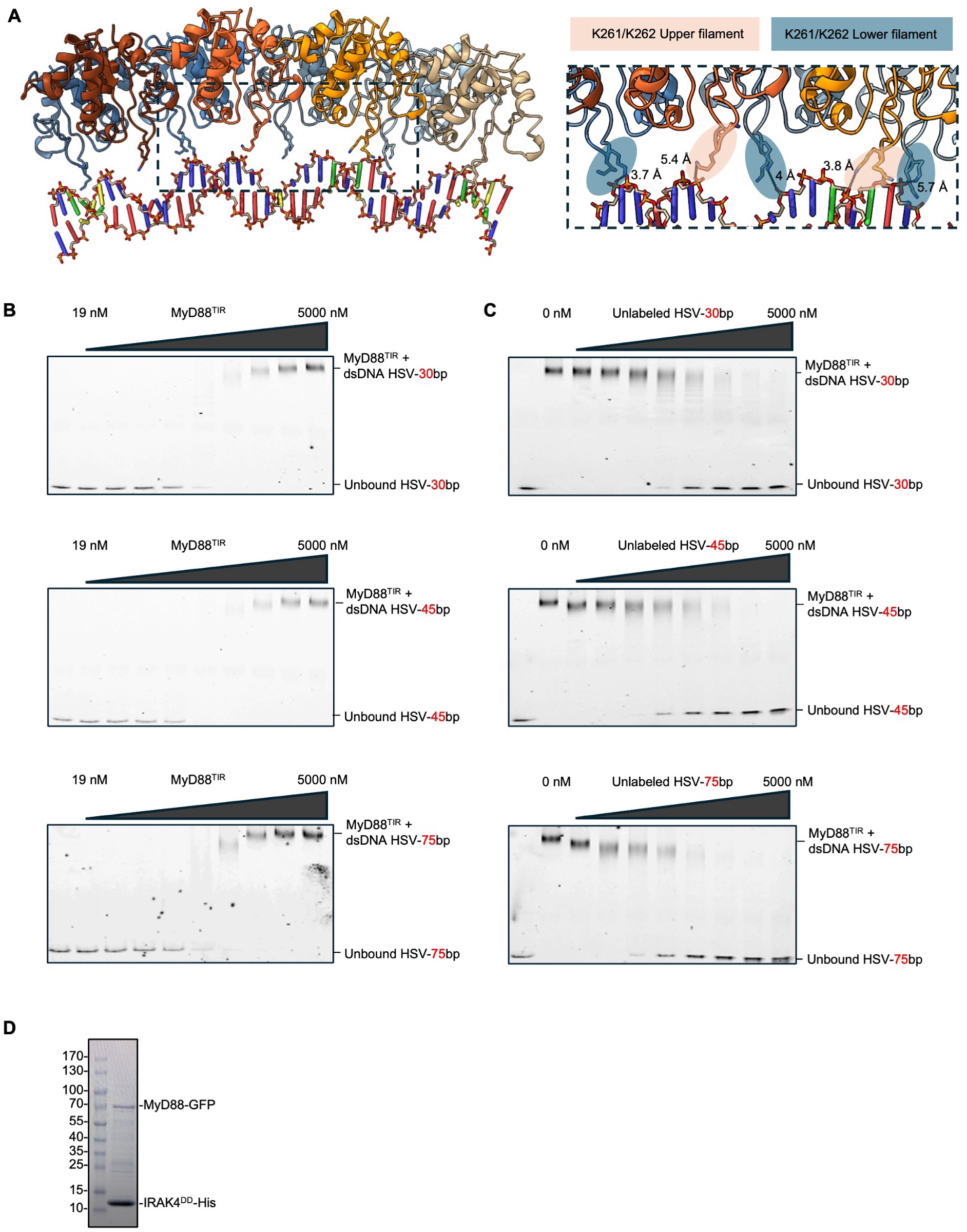
Electrophoretic mobility shift assays (EMSAs) of MyD88^TIR^. **(A)** Detailed interaction of WT MyD88 with dsDNA (left) and a close-up view (right) of lysine residues K261 and K262 positioned near the negatively charged phosphate backbone of the dsDNA. Distances between several K261 and K262 residues and phosphate backbone phosphoryl oxygen atoms are shown in Å. **(B)** EMSAs of increasing amount of MyD88^TIR^ with dsDNA HSV-30bp, HSV-45bp, and HSV-75bp. MyD88^TIR^ concentrations: 19, 39, 78, 156, 312, 625, 1,250, 2,500 and 5,000 nM. Cy3 dsDNA: 0.5 nM. Fitted KD and Hill coefficient are reported in Figure 2. **(C)** Competition EMSAs of MyD88^TIR^ with HSV-30bp, HSV-45bp and HSV-75bp dsDNA using increasing amount of unlabelled DNA. MyD88^TIR^ concentration: 2,000 nM. Cy3 dsDNA: 0.5 nM. Unlabelled DNA: 8, 20, 51, 128, 320, 800, 2,000, 5,000 nM. **(D)** Full-length MyD88-GFP was co-expressed with His-tagged IRAK4 DD in E. coli BL21 (DE3). The complex was co-purified by nickel affinity chromatography and analysed by SDS-PAGE.

**Figure S7.**
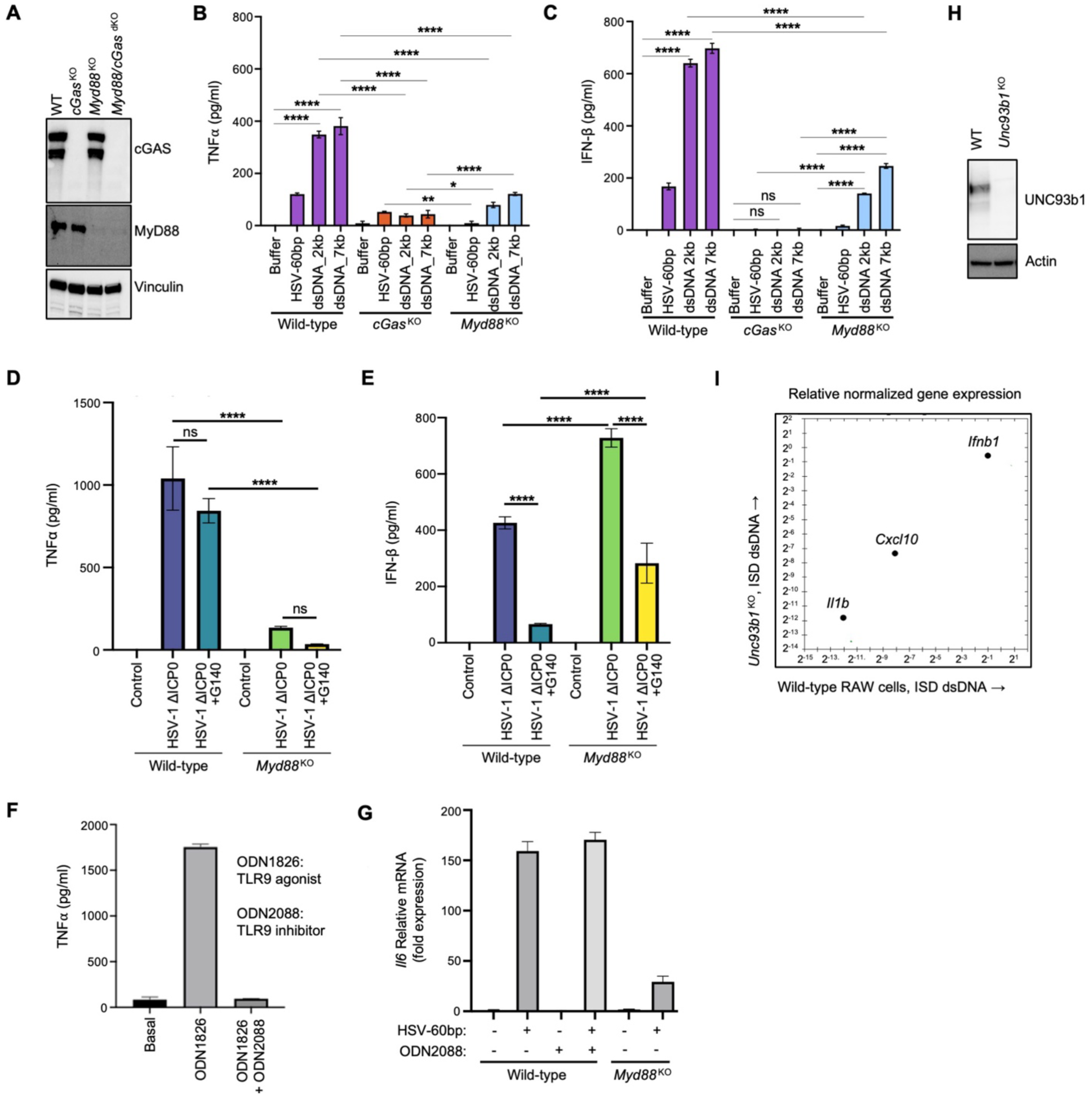
MyD88 and cGAS in dsDNA-induced cytokine release and signalling. **(A)** Verification of *cGas* KO, *Myd88* KO, and *cGas/Myd88* dKO by western blots. **(B)** TNFα release analysed by ELISA. WT, *Myd88* KO or *cGas* KO iBMDMs were transfected with either short HSV-1 dsDNA (60 bp) or long dsDNA fragments (2.5 µg) representing the length boundaries of 2,000 bp and 7,000 bp, respectively, for 6 h. **(C)** IFN-β release analysed by ELISA. WT, *Myd88* KO or *cGas* KO iBMDMs were transfected with either short HSV-1 dsDNA (60 bp) or long dsDNA fragments (2.5 µg) representing the length boundaries of 2,000 bp and 7,000 bp, respectively, for 6 h. **(D, E)** TNFα (D) or IFN-β (E) release analysed by ELISA. WT or *Myd88* KO primary BMDMs were infected with HSV-1 ΔICP0 strain for 18 h. The indicated cells were pretreated with G140 (10 µM) for 12 h. Data in bar graphs are represented as mean ± SEM. The designation of *, **, *** and **** correspond to p ≤ 0.05, p ≤ 0.01, p ≤ 0.001, and p ≤ 0.0001, respectively. Conditions were compared by using Ordinary One-way ANOVA. **(F)** Pretreatment of TLR9 inhibitor ODN2088 (1 µM) for 15 min inhibited TNFα secretion by the TLR9 agonist ODN1826 (1 µM) treated for 24 h. **(G)** Fold induction of *Il6* mRNA expression analysed by qRT-PCR upon transfection of HSV-60bp dsDNA (2.5 µg) for 6 h. *Il6* mRNA upregulation was not affected by pretreatment of TLR9 inhibitor ODN2088 (1 µM) for 15 min. **(H)** Verification of *Unc93b1* KO RAW 264.7 macrophages by western blots. **(I)** Relative mRNA expression of Il1b, Cxcl10, and Ifnb1 determined by qRT-PCR. WT or *Unc93b1* KO RAW 264.7 macrophages were transfected with inflammatory stimulatory dsDNA (2.5 µg) for 6 h.

**Figure S8.**
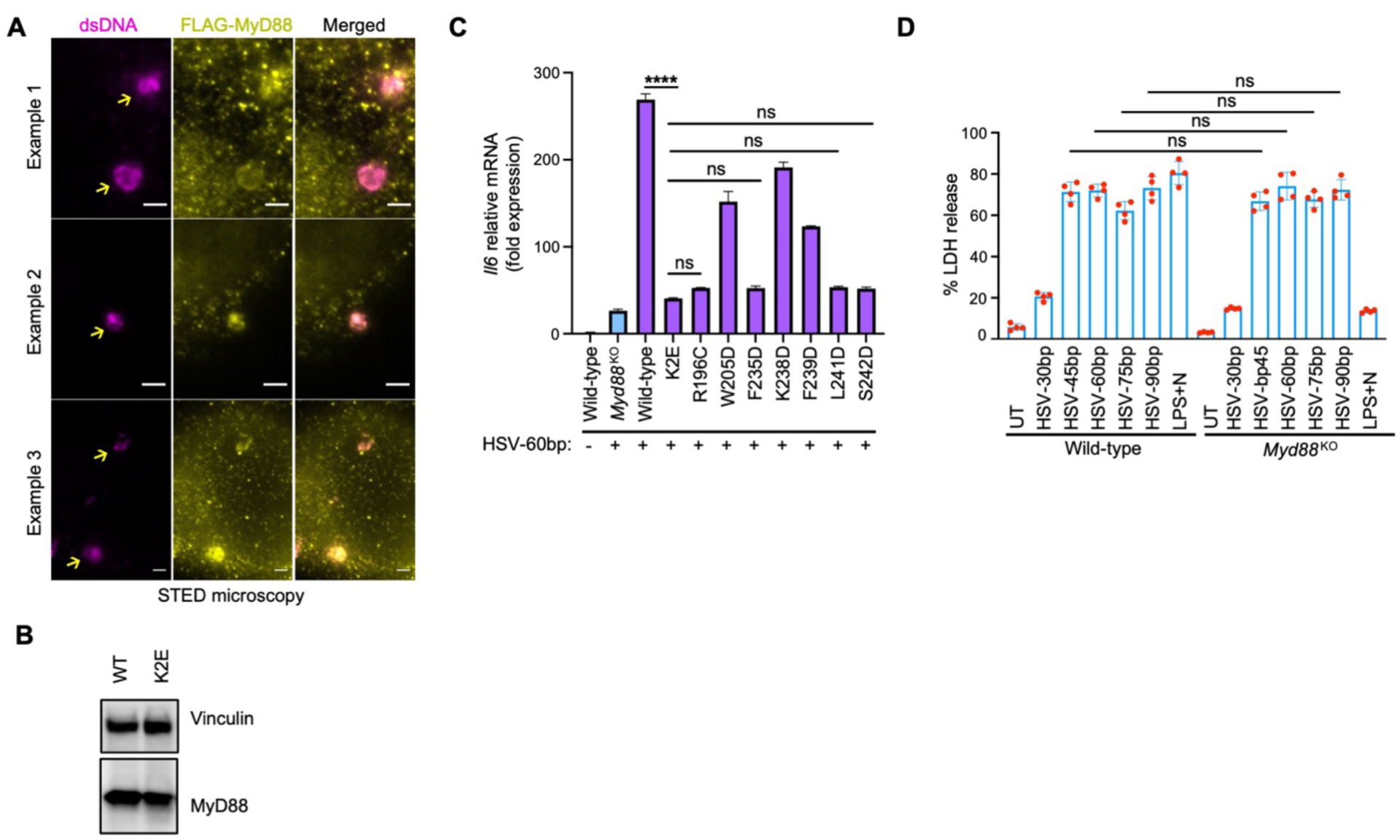
MyD88 is a sensor for dsDNA-mediated NF-κB activation. **(A)** Three examples of stimulated emission depletion microscopy (STED) imaging. WT iBMDMs with *Myd88* endogenously tagged by GFP and FLAG were transfected with HSV-60bp. Fixed cells were co-stained with anti-dsDNA and anti-FLAG antibodies. Puncta of dsDNA and MyD88 in approximate barrel shapes are labelled by arrows. The images are representative of n=15 experiments. Scale, 1 µm. **(B)** Western blot of MyD88 in *Myd88* KO iBMDMs reconstituted with WT or K2E. **(C)** WT, *Myd88* KO, *Myd88* KO reconstituted with WT, K2E, or mutations at the interstrand interface (R196C, W205D, F235D, K238D, F239D, L241D, S242D) were transfected with HSV-60bp dsDNA (2.5 µg) for 6 h. Fold change of *Il6* mRNA was determined by qRT-PCR. **(D)** WT or *Myd88* KO iBMDMs were transfected with the indicated lengths of HSV-1 dsDNA (2.5 µg) for 6 h. The indicated cells were primed with LPS (1 μg/ml) for 4 h and treated with nigericin (20 μM) for 30 minutes. LDH release was monitored and normalized to maximum LDH content of each cell line. Data in bar graphs are represented as mean ± SEM. The designation of *, **, *** and **** correspond to p ≤ 0.05, p ≤ 0.01, p ≤ 0.001, and p ≤ 0.0001, respectively. Conditions were compared by using Ordinary One-way ANOVA.

**Figure S9.**
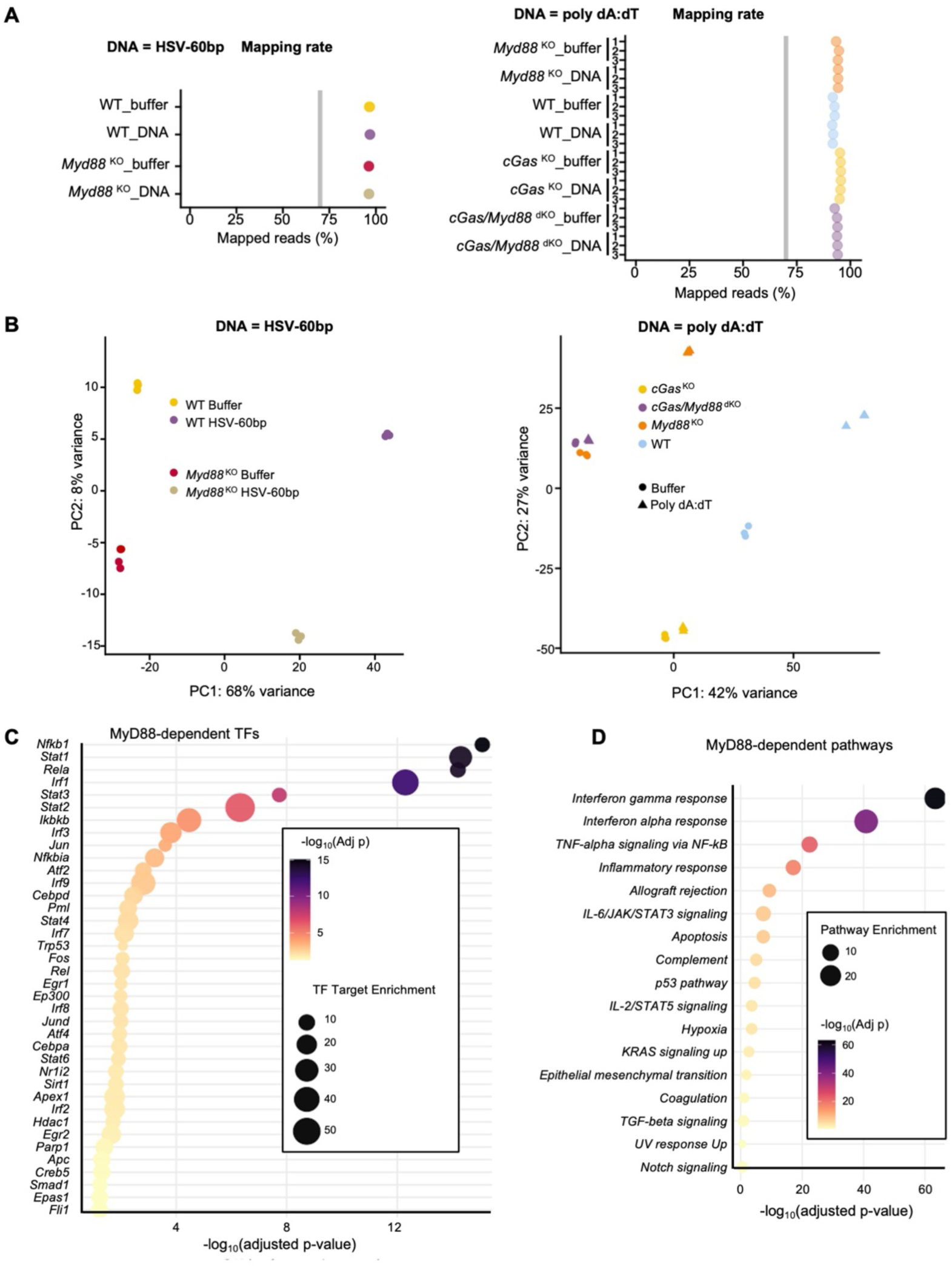
RNA-seq reveals the role of MyD88 in cytosolic DNA-induced transcriptional activation. **(A)** Assessment of 5′ to 3′ bias across all samples for the 2 RNA-seq datasets. The plots depict the 5′ to 3′ coverage bias values for each experimental condition, with values close to 1.00 indicating minimal bias and consistent transcript coverage along the gene body. **(B)** Principal Component Analysis (PCA) of all samples. Each plot displays the distribution of samples along the first two principal components (PC1 and PC2), illustrating the variance and clustering based on global gene expression profiles. Samples clustered together exhibit similar expression patterns. **(C, D)** Transcription factor (C) and pathway (D) analyses from genes downregulated by *Myd88* KO under HSV-60bp transfection.

**Figure S10.**
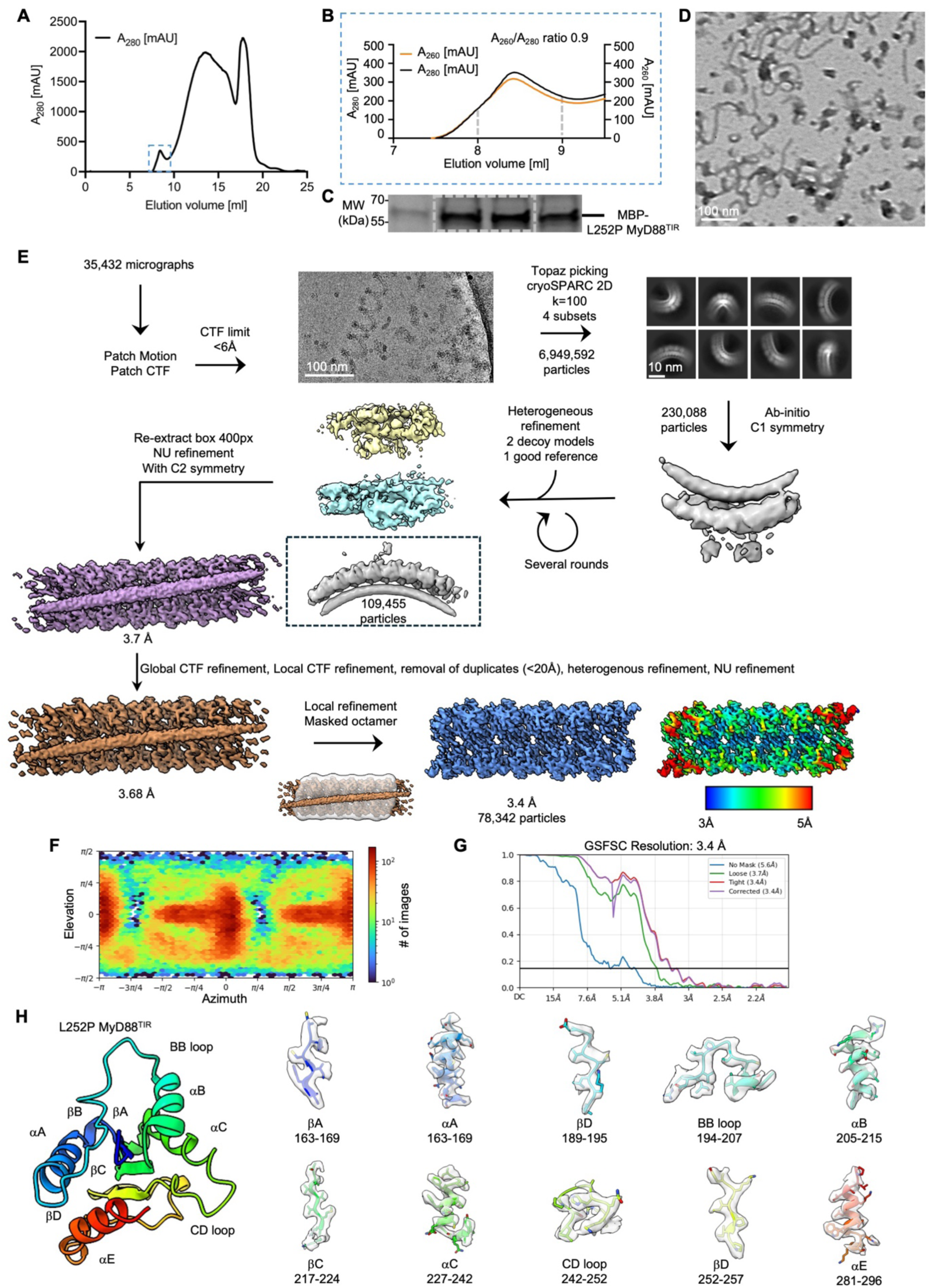
Cryo-EM processing workflow of the L252P MyD88^TIR^ spiral segment. **(A, B)** Size-exclusion chromatography profiles from the final purification step of the L252P MyD88^TIR^ sample, with the complete profile (A) and an enlarged view of the void volume (B). **(C)** SDS-PAGE analysis with Coomassie staining of fractions 7.5 to 9.5 ml from (B). **(D)** Negative stain EM analysis of the L252P MyD88^TIR^ sample from the void volume. Scale bar, 100 nm. **(E)** Data processing workflow for the L252P MyD88^TIR^ spiral, detailing stages from initial micrograph collection to 2D classification and final local refinement. Scale bars: micrograph, 100 nm; 2D class averages, 10 nm. **(F, G)** Orientation diagnostics from the final local refinement step, showing distribution of particle orientations (F) and the Fourier shell correlation (FSC) plot (G). **(H)** Map quality evaluation of the L252P MyD88^TIR^, the cryo-EM map regions are displayed as transparent surface representations, with corresponding atomic models fitted into the density and shown in cartoon representation to highlight structural details.

**Table S1. Cryo-EM data collection, refinement and model validation Statistics.**

**Table S2. RNA-seq results showing differential gene expression analyses in WT and *Myd88* KO iBMDMs treated with HSV-60bp DNA.**

**Table S3. RNA-seq results showing differential gene expression analyses in WT, *Myd88* KO, *cGAS* KO, and *Myd88/cGAS* dKO iBMDMs treated with polydAdT DNA.**

**Video S1. Overall structure of the MyD88 TIR domain ring segment.**

**Video S2**. **A morph demonstrating the flexibility of bound dsDNA and the MyD88 TIR domain segment as captured by 3D classification**. The video reveals subtle conformational adjustments in both the DNA and MyD88 TIR domain.

**Video S3**. **A morph video illustrating structural differences in the MyD88^TIR^ protomer between the WT and L252P models**. The video begins with the WT structure and transitions to the L252P TIR domain model, highlighting the conserved nature of the overall protein structure. Notably, the CD loop emerges as the primary region exhibiting structural differences.

**Video S4**. **This video of the central MyD88^TIR^ segment octamer shows the global structural changes caused by the L252P mutation**. The video starts with the WT model and then morphs to the L252P octamer structure. The morph is shown in different orientations to highlight how the local conformational change in the CD loop is translated into global structural changes.

